# Plasma electrolyte changes can modulate post-repolarisation refractoriness in the human atrium

**DOI:** 10.64898/2026.11.20.768085

**Authors:** Cesare Corrado, Caroline H. Roney, Sanjiv M. Narayan, Wayne R. Giles, Steven A. Niederer

## Abstract

Changes in plasma Na^+^, K^+^ and Ca^2+^ levels are common in patients with chronic kidney disease or gastrointestinal disorders, or in those taking certain medications, and may promote atrial fibrillation or impair drug-induced cardioversion. Post-repolarisation refractoriness (PRR) is an established mechanism by which antiarrhythmic drugs suppress tachyarrhythmias. We investigated how changes in extracellular Na^+^, K^+^ and Ca^2+^ concentrations ([Na^+^]_o_, [K^+^]_o_, [Ca^2+^]_o_) modulate ionic currents and PRR in healthy and fibrotic human atrial tissue. We used a multi-scale computational framework comprising an isolated atrial myocyte, a one-dimensional atrial tissue strand, and a virtual cohort of 14 left atrial digital twins, combined with Gaussian process emulators and variance-based global sensitivity analysis. Changes in [K^+^]_o_ strongly modulated PRR by altering the inward rectifier K^+^ current (IK1), shifting the resting potential and thereby regulating Na^+^-current availability. In contrast, [Na^+^]_o_ had only a weak effect on PRR, mediated primarily through the Na^+^/Ca^2+^ exchanger, and [Ca^2+^]_o_ had a negligible effect. Tissue PRR was more sensitive to electrolyte changes than myocyte PRR, reflecting electrolyte-dependent alterations in the safety factor for action potential propagation. Changes in [K^+^]_o_ also substantially affected arrhythmia vulnerability, increasing inducibility and prolonging rhythm disturbances. We conclude that extracellular K^+^ is the principal plasma electrolyte determinant of atrial PRR and arrhythmia burden, acting primarily through IK1. Monitoring and maintaining serum K^+^ within the normal range may be important for managing atrial arrhythmia risk.

**Author summary:** Atrial fibrillation is the most common heart rhythm disorder and increases the risk of stroke, heart failure and death. Many people with this condition also have abnormal levels of electrolytes, such as potassium, sodium and calcium, in their blood. These changes are common in people with kidney or digestive disorders and in those taking certain medications. We wanted to understand how changes in these electrolytes affect post-repolarisation refractoriness (PRR), a protective property of heart tissue that prevents it from being excited again for a short time after each heartbeat. Some anti-arrhythmic drugs work by prolonging this period. Since it is difficult to study the effects of individual electrolytes in patients, we used computer models, ranging from single heart cells to patient-specific digital twins of the atria from 14 patients, together with statistical models that predict their behaviour. We found that potassium had by far the strongest effect on PRR, acting mainly through a specific potassium channel. Its influence was also greater in whole heart tissue than in individual cells. Abnormal potassium levels made arrhythmias easier to trigger and more likely to persist. Our findings highlight the importance of blood potassium levels in managing the risk of atrial arrhythmias.

## 1 Introduction

Atrial fibrillation (AF) is the most common sustained human cardiac arrhythmia, with a recently estimated prevalence of 33.5 million cases worldwide [1] and an increasing trend, partly due to population ageing [2]. AF is characterised by very rapid rates and disorganised rhythms [3]. Importantly, AF can be a significant cause of stroke, heart failure and death [4].

Slow conduction velocity (CV), combined with relatively short effective refractory periods (ERP), are known to provide a favourable substrate to initiate and maintain AF in the human atrium [5,6]. Alterations in plasma electrolytes occur frequently and can significantly modify both ERP and CV. For example, approximately 20% of patients admitted for cardioversion present with hypokalaemia [7]. The incidence of hyperkalaemia is smaller; nevertheless, it is also a clinically important challenge [8]. Epidemiological evidence also links higher sodium intake (but not potassium intake alone) with increased AF prevalence [9].

Previous studies have revealed that alterations in atrial refractoriness cannot be fully accounted for by changing the action potential duration (APD). Rather, the atrial myocyte may repolarise completely and yet remain unexcitable. Thus, its ERP outlasts repolarisation. This difference has been denoted post-repolarisation refractoriness (PRR), and it is defined as the difference between the ERP and the APD measured at 75% of full or complete repolarisation [10]. The resting membrane potential and the availability of the fast Na^+^ current (INa) [11,12] primarily govern PRR in mammalian hearts. Both factors can be modulated by the plasma concentration of potassium, [K^+^]_o_, and their functional linkage. This is mainly due to changes in the nonlinear inward rectifier K^+^ current (IK1) [13].

Given its incidence and pathophysiological significance, improved approaches for clinical management of AF are being actively investigated. It has been observed that raising [K^+^]_o_ can convert chronic AF to sinus rhythm [14]. Moreover, changes in [K^+^]_o_ can potentiate the effects of Na^+^-channel blockers [15] in the setting of AF. Even small changes in PRR are now a recognised antiarrhythmic mechanism in the human heart [16], with the Na^+^-channel-blocking drugs (e.g. pilsicainide [17], ranolazine [18]) significantly enhancing PRR values.

The main goal of this study was to determine how clinically relevant variations in extracellular electrolytes, [Na^+^]_o_, [K^+^]_o_, or [Ca^2+^]_o_, can alter human atrial PRR and contribute to anti-arrhythmia endpoints. However, determining how plasma electrolytes can independently alter PRR is difficult both in vitro and in vivo. Accordingly, we have used biophysically detailed human atrial electrophysiological myocyte models [19], that have been demonstrated to be effective in studying in silico the effects on an isolated myocyte of variations in [Ca^2+^]_o_ [20] and low [K^+^]_o_ [21]. In a previous study [22], we used left atrial digital twins to study how mild variations in [Na^+^]_o_, [K^+^]_o_ and [Ca^2+^]_o_ modulate the onset (inducibility) and the sustainability of human atrial arrhythmias.

Specifically, in this study, we quantify in silico how clinically relevant variations in [K^+^]_o_, [Na^+^]_o_ and/or [Ca^2+^]_o_ can alter atrial PRR. We also identify the changes in the transmembrane currents that produce these alterations in (i) the isolated myocyte, as well as (ii) a strand of atrial tissue. These assessments are done in both healthy (baseline) conditions and in the settings of atrial fibrosis. The resulting electrophysiological insights provide the basis for using personalised human atrial data sets to analyse how these plasma electrolyte alterations can affect the inducibility and duration of the atrial arrhythmias. Our multiscale computational approach (an isolated human atrial myocyte, a one-dimensional tissue strand, and a cohort of patient-specific left atrial digital twins [23,24]) also employs global sensitivity analysis [25]. Gaussian process emulators [26–28] are utilised to manage the numerical complexity arising from the computational cost of simulating atrial arrhythmias.

## 2 Materials and methods

### 2.1 The human atrial myocyte model

The Courtemanche [19] model of the human atrial action potential (AP) accurately simulates most electrophysiological properties of this myocyte. However, as we have done previously, modified expressions for the delayed rectifier K^+^ current (IKr) and the inward rectifier K^+^ current (IK1) were incorporated in this study to provide an accurate reproduction of the variations of the APD and the resting potential when [K^+^]_o_ is altered [22].

In addition, the expression for the L-type calcium current (ICaL) was modified so that it included a variable reversal potential. This was based on the standard Nernst equation:

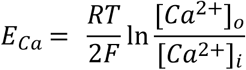

The initial baseline electrolyte concentrations were set as follows: [Na^+^]_o_ = 140 mM; [K^+^]_o_ = 4.5 mM; [Ca^2+^]_o_ = 1.16 mM, according to reported values for human plasma [20,22].

Atrial fibrosis was modelled using the same approach that we and others have previously adopted [22]. In brief, this consists of reducing the fast inward sodium current (INa) in the atrial myocyte to 60% of its baseline values, and reducing both the ICaL and IK1 in the atrial myocyte by 50%.

### 2.2 Clinically relevant variations in plasma electrolyte concentrations

Based in part on our previous study [22], we selected the following ranges for 3 different electrolyte variations: [Na^+^]_o_ 105-175 mM, which covers severe hyponatraemia ([Na^+^]_o_≤125 mM) to severe hypernatraemia ([Na^+^]_o_≥160 mM); and [K^+^]_o_ was varied between 2 and 10 mM, thus accounting for both severe hypokalaemia ([K^+^]_o_≤2.5 mM) and severe hyperkalaemia ([K^+^]_o_≥6.5 mM). This range is also known to result in biphasic behaviour of conduction velocity during hyperkalaemia (this phenomenon appears when [K^+^]_o_≥8 mM). [Ca^2+^]_o_ was varied in the range [0.5-2.5] mM, which spans severe hypocalcaemia ([Ca^2+^]_o_≤0.5 mM) and significant hypercalcaemia ([Ca^2+^]_o_≥2.5 mM).

### 2.3 Protocol to evaluate post-repolarisation refractoriness in the human atrial myocyte

According to previous reports, post-repolarisation refractoriness (PRR) was defined as the difference between the effective refractory period (ERP) and the APD measured at 75% of full repolarisation (APD_75_) [10]. In this study, most PRR values were obtained at four different cycle lengths (CL): 1000, 500, 400 and 300 ms. The following procedure was used: (i) The diastolic threshold (DI_tsh_) for AP excitation was operationally defined as the smallest intensity of a stimulus (with a duration of 2 ms) that elicits an AP after a long period of quiescence (independent of the cycle length). These DI_tsh_ values were obtained with a resolution of 1 pA/pF. In all subsequent parts of this study, a stimulus intensity 3 times DI_tsh_ was applied. (ii) Each atrial myocyte was paced for 15 minutes to reach a limit cycle [21], and the APD_75_ was computed using data from each of the last two action potential traces (APD_75,1_, APD_75,2_) and the resting membrane potential (RMP) (RMP_1_, RMP_2_). For each of these two AP traces, we also computed the following quantities: APD_90_; action potential amplitude, ΔV, between the RMP and the plateau; the maximum upstroke velocity 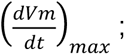 the area under the action potential trace (AUAPT). (iii) The corresponding effective refractory periods (ERP_1_, ERP_2_) were obtained by applying a premature stimulus S2 (with 1 ms resolution) and marking as ERP the largest S2 that did not produce a regenerative AP. Each ERP was estimated using a bisection search method within the range [max(0.7APD_90_, 50 ms), CL ]. When APD_90,1_ and APD_90,2_ differed by less than 5% (that is, no APD alternans occurred), we computed one ERP only. Alternatively, in the presence of APD alternans, we computed one ERP for each of the two action potentials that exhibited alternans. Note that we did not include any ERP values when the atrial myocyte exhibited a 2:1 activation pattern.

To account for and systematically deal with stimulus artefact (which tended to be prominent after the larger variations in plasma electrolytes, and in the settings of fibrosis), we defined stimulus capture when all the following conditions are met:

- APD_90_^S2^ ≥ 50 ms
- ΔV^S2^ ≥ max (30 mV, 0.5 ΔV^S1^)
- 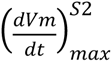≥ max(30 V/s, 0.2 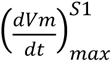)
- AUAPT^S2^ ≥ 0.5 AUAPT^S1^
- min _tɛ[0,5] ms_ (I) <0

where I_ion_ represents the total transmembrane ionic current generated by the atrial myocyte, and the superscripts S1, S2 indicate the quantities evaluated at the selected cycle length (S1) and in the presence of a premature stimulus (S2).

### 2.4 Protocol to evaluate post-repolarisation refractoriness and conduction velocity in the human atrial tissue strand

In addition to PRR assessment using a single human atrial myocyte, we also studied electrolyte-induced changes in PRR and CV in human atrial tissue. This was done (as diagrammed in Fig 1) using a 10 mm long homogeneous rectangular atrial tissue strand having a cross-sectional dimension of 100 μm × 100 μm. This atrial strand was discretised by tetrahedral elements with 100 μm edges, generated using the mesher utility from openCARP software [29]. At baseline, we adopted a conductivity of 0.4 S/m, yielding a conduction velocity of approximately 0.32 m/s [30] in the presence of reference electrolyte concentrations and a stimulus CL of 1000 ms (1 Hz). To simulate the setting of fibrosis, the conductivity was set to 0.31 S/m, and the CV was 0.24 m/s. A monodomain description was adopted, in which cardiac tissue is represented as a single continuous electrical medium rather than modelling the intracellular and extracellular spaces separately [31]. PRR values were computed for cycle lengths of 1000, 500, 400 and 300 ms. For each selected cycle length, the myocytes were paced for 15 minutes to reach a limit cycle [21]. We applied two consecutive stimuli of amplitude 60 pA/pF and a duration of 1 ms over the first 400 μm of the left end of the atrial strand. The corresponding electrophysiological responses were recorded from the start of these two APs to the end of a 1000 ms safety window (total window: 2 × CL + 1000 ms). The ‘tissue stat’ was saved at t = 2 × CL for subsequent S2 simulations. Time discretisation was set at dt = 10 μs, and the solution was sampled every dt_out_ = 100 μs.

**Fig 1.**
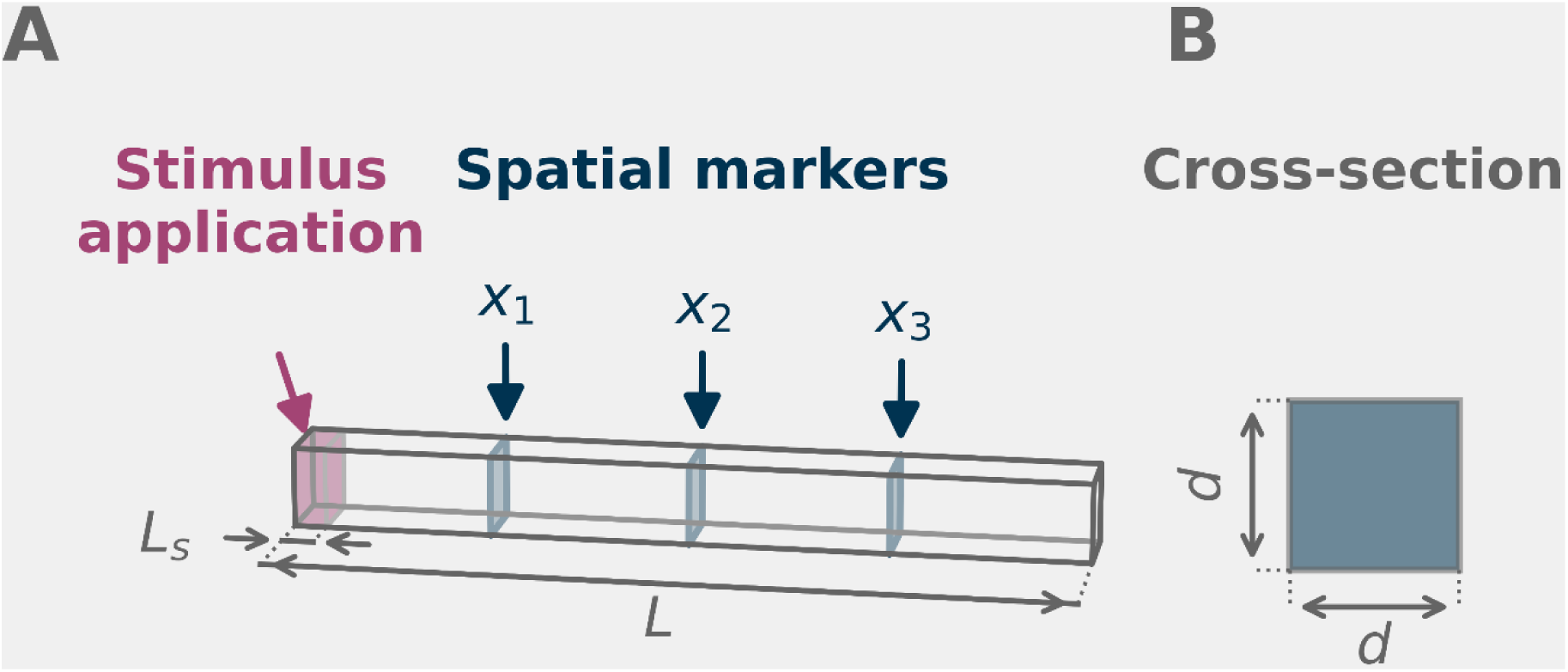
Diagram of the atrial tissue strand showing its dimensions and the stimulus and measurement details. An atrial tissue strand of length L=10 mm (Panel A) with a square cross-section of side d=100 μm (Panel B) was used to measure conduction velocity and action potential duration. The stimulus was applied on the left end, over a length L_s_ =400 μm. The three shaded planes in Panel A are the spatial markers x_1_ = 2.5 mm, x_2_ = 5 mm and x_3_ = 7.5 mm, measured from the stimulated end. APD was measured in the central section x_2_. CV was measured as the mean velocity of the activation front between x_1_ and x_3_.

At the spatial markers x_1_=2.5 mm, x_2_=5 mm and x_3_=7.5 mm (see Fig 1), the three traces of the transmembrane potential over each of the cross-sections were extracted. We evaluated APD_75_ at x_2_ = 5 mm, and the mean CV between x_1_=2.5 mm and x_3_=7.5 mm for each of the two final consecutive S1-generated action potentials elicited by the stimulus train.

Activation times at each selected spatial location were detected as 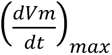 using a 50 ms search window which started at the estimated wave-front arrival time (estimated from the propagation delay between adjacent sites). The procedure described for evaluating excitability/refractoriness in the isolated atrial myocyte was repeated when in silico atrial strands were utilised. Successful stimulus capture required the following 3 conditions to be met:

- At x_1_=2.5 mm, x_2_=5 mm, and x_3_=7.5 mm, the amplitude of the action potential is larger than 10 mV
- At x_1_=2.5 mm, x_2_=5 mm, and x_3_=7.5 mm 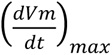 > 10 *V*/*s*
- The mean CV between x_1_=2.5 mm and x_3_=7.5 mm is larger than 0.01 m/s

Each ERP value was obtained using a bisection search method, with search interval minimum equal to 70% of the APD_90_ measured at x=2.5 mm (or 50 ms, whichever is larger) and search interval maximum equal to the cycle length.

### 2.5 Global sensitivity analysis

An established variance-based method [25] was applied to data obtained at each stimulus CL to perform the global sensitivity analysis (GSA) of PRR values in response to selected variations of plasma electrolyte concentrations. GSA was also employed to estimate the extent to which individual transmembrane ionic currents can modulate PRR. Briefly, this approach expresses the output variance of the PRR values as a sum of contributions from changes in the input parameters (electrolyte concentration values), and can also capture their interactions. First-order effects measure the direct impact on the output when each input parameter is varied while keeping the other input parameters fixed. Second- and higher-order effects capture the interactions between two or more selected input variables, thus revealing their combined impact on changes in PRR. Finally, the ‘total effects’ express the overall influence of each selected input parameter on PRR. This total variance accounts for the first-order effect as well as the effects due to interactions with all other variables. These changes in variance are expressed using Sobol indices, defined as the ratio between the variance of the selected effect (first-, second- and higher-order) and the total variance. We computed Sobol indices using SALib [32] which is based on numerical integration with N(2D + 2) points (samples) from the input parameter space, with D being the dimension of the input space and N a user-defined hyperparameter. In this study, we chose N=2048.

To make this analysis computationally tractable, we have adopted a surrogate model consisting of a simplified, computationally efficient approximation of the simulated procedure. Specifically, we utilised the Gaussian process (GP) provided by GPErks (available at https://pypi.org/project/GPErks). To obtain the GSA of the PRR in response to the selected electrolyte variations, we fitted the GP over training sets of 300 Latin hypercube samples (LHS). To obtain the GSA in response to each change in the ionic currents that underlie the atrial action potential, we trained the GP using a training data set of 40 × N_channels_ (myocytes) and 100 × N_channels_ (tissue) LHS samples. Here, N_channels_ represents the number of ion channels selected for analysis.

In this study, we estimated the sensitivity of PRR to each transmembrane ion current by first identifying which of these conductances were sensitive to alterations in a specific plasma electrolyte. In the case of (i) Na^+^-sensitive channels, we studied INa, IbNa, INaK and INaCa (4 currents); (ii) for K^+^-sensitive currents, we focused on IK1, IKr, IKs, Ito, IKur and INaK (6 currents); and (iii) for Ca^2+^-sensitive channels, we studied INaCa, ICaL and IbCa (3 currents). For example, to estimate the sensitivity to Ca^2+^-modulated currents, we ‘applied’ each selected change in [Ca^2+^]_o_ to [Ca^2+^]_o, INaCa_, [Ca^2+^]_o, ICaL_, and [Ca^2+^]_o, IbCa_ to ensure that each contribution was varied independently of the others.

When studying the sensitivity of the human atrial arrhythmias to variations in plasma electrolytes, we restricted our analysis to variations in [Na^+^]_o_ and [K^+^]_o_. This is because PRR and CV showed a negligible sensitivity to variations in [Ca^2+^]_o_ in the modelling approach that we have utilised. A Latin hypercube design was utilised to generate a matrix of 40 ([Na^+^]_o_, [K^+^]_o_) pairs of these two plasma electrolyte concentrations, and each of these was applied to the entire digital twin cohort.

### 2.6 Human left atrial digital twins

An important part of this study is based on simulated induced atrial arrhythmias done by using a data set comprised of 14 clinical cases adapted from a previously published human atria data repository [24]. All patient data were previously published and fully anonymised; ethical approval and informed consent were obtained in the original studies [24]. The selection criteria are described in S1 Appendix, Section A2. Each patient data set consists of detailed entire left atrial anatomical parametrisations (triangulations) supplemented by image intensity ratio (IIR) maps, fibre directions and universal atrial coordinates (UAC) [23].

To obtain a computational mesh suitable for numerical simulations, we refined the triangulations describing the atrial anatomy [24] to obtain a maximum edge length of 440 µm, using the adaptive subdivision algorithm implemented in the VTK [33] library.

For each clinical case, we separated portions of the atrial tissue into 6 regions or subtypes, based on their IIR values. The six columns in Table 1 summarise the IIR ranges used to identify each subtype or region. The 5 rows in Table 1 report the parameter values assigned to each subtype. These correspond to the longitudinal and transverse conductivities (σ_il_, σ_it_), and the conductances of the fast inward Na^+^ current (GNa), the inward-rectifier K^+^ current (GK1) and the L-type Ca^2+^ current (GCaL) that were assigned in our approach for simulating atrial fibrosis [34].

**Table 1.** Human left atrial tissue electrophysiological parameters (column 1), expressed as a function of their value of image intensity ratios (IIR; columns 2-7).

|  | **IIR ≤ 0.9** | **0.9 < IIR ≤ 1.1** | **1.1 < IIR ≤ 1.22** | **1.22 < IIR ≤ 1.4** | **1.4 < IIR ≤ 1.6** | **IIR > 1.6** |
|---|---|---|---|---|---|---|
| **σ<sub>il</sub> (S/m)** | 0.4 | 0.31 | 0.31 | 0.31 | 0.28 | 0.19 |
| **σ<sub>it</sub> (S/m)** | 0.1 | 0.075 | 0.075 | 0.075 | 0.07 | 0.0475 |
| **GNa (nS/pF)** | 7.8 | 7.8 | 7.8 | 4.68 | 4.68 | 4.68 |
| **GK1 (nS/pF)** | 0.09 | 0.09 | 0.09 | 0.045 | 0.045 | 0.045 |
| **GCaL (nS/pF)** | 0.1238 | 0.1238 | 0.1238 | 0.0619 | 0.0619 | 0.0619 |

### 2.7 Simulation of induced atrial arrhythmias and their endpoint classifications

Each atrial arrhythmia was initiated by utilising three different stimulus protocols. In the first two, the following procedure was adopted [35]. The Courtemanche human atrial model was initialised in each of the six designated left atrial regions by pacing it for 15 minutes at a CL of 700 ms [21]. Following this, an arrhythmia induction protocol was immediately applied. This consisted of 5 consecutive stimuli applied at a CL of 700 ms in the proximity of the coronary sinus (CS), followed by a 400 ms ‘blanking period’, and then a short burst-pacing challenge consisting of a train of 5 stimuli delivered sequentially at a CL of 160 ms in two different ‘ectopic’ locations (protocols 1 and 2). These locations were in the proximity of (i) the left superior pulmonary vein (LSPV) or (ii) the right superior pulmonary vein (RSPV). Both of these targeted pacing regions were considered to be circular, with a radius of 2 mm. The centres of each and of CS was localised using UAC. The UAC ranges (minimum and maximum of each component), containing each designated centre, were manually identified on one randomly chosen human atrial dataset:

- CS within 0.5≤UAC1≤0.7; 0.8≤UAC2≤0.9,
- LSPV within 0.86≤UAC1≤0.88; 0.65≤UAC2≤0.75,
- RSPV within 0.05≤UAC1≤0.08; 0.6≤UAC2≤0.63.

For each patient data set and for each pacing location, the 3D space coordinates of the points that mapped to the previously defined UAC ranges were identified and used to compute the barycentre. This point was then used as the centre where pacing stimuli were delivered.

In contrast, in the third protocol, each left atrial tissue was first paced for 15 minutes at a cycle length of 160 ms [36], and then an initial ‘arrhythmia induction condition’ consisting of isochrones of 4 spiral voltage waves [37] was immediately applied. These isochrones were computed using the UAC as previously described [23,36].

In this study, for each human atrial data set or ‘preparation’, all three arrhythmia induction protocols were applied using openCARP software [29]. A time step Δt=5 μs was used, and each simulated transmembrane potential trace was sampled every 1 ms and saved. Each stimulus-induced atrial arrhythmia was recorded for a total of 10 seconds. This allowed us to obtain descriptive parameters that are consistent with analogous data sets published by other groups [38]. For each, we also computed the duration of the arrhythmia (T_term_) using the procedure described in S1 Appendix, Section A3.

We observed that the simultaneous presence of severe hyperkalaemia and hypernatraemia consistently produced markedly slowed conduction within fibrotic atrial tissue, thus creating a large conduction velocity gradient between healthy and fibrotic regions. That is, under these conditions, activation propagated very slowly through the fibrotic substrate before rapidly traversing the surrounding healthy tissue and re-entering the fibrotic region. These factors also resulted in functional macro-re-entrant circuits having long cycle lengths. Consequently, simulations with T_term_ between 5000 and 9900 ms were visually inspected. Based on this inspection, simulations with a termination time T_term_ ≥8900 ms consistently exhibited sustained re-entry, whereas those with shorter durations represented self-terminating activity. Therefore, arrhythmias were classified as self-sustained only when T_term_ ≥8900 ms, and the organisation index (OI) was computed following the procedure described in S1 Appendix, Section A4.

In the final step of this arrhythmia classification exercise, each simulated output was ‘placed’ into one of the following 4 endpoints:

- non-initiated arrhythmia: that is, an absence of atrial substrate activations after the induction protocol;
- self-terminating atrial arrhythmias: i.e. those lasting less than 8900 ms;
- sustained atrial tachycardia (AT): i.e. lasting ≥ 8900 ms with OI>0.6;
- sustained atrial fibrillation (AF): i.e. lasting ≥ 8900 ms with OI≤0.6.

We note that the threshold OI=0.6 used to discriminate between AT and AF corresponds to the largest OI that has been recorded in the proximity of the coronary sinus during clinical procedures in patients presenting with AF [39,40]. These endpoints were ordered based on ‘severity’, with sustained arrhythmias defined as the most severe (AF more severe than AT), followed by self-terminating arrhythmias and finally non-inducibility. For each patient and each plasma electrolyte concentration pair, this analysis was completed by selecting and designating the most severe endpoint exhibited by the data set in response to the 3 induction protocols. We adopted a similar principle for the duration of the arrhythmia (T_term_), selecting and designating the longest T_term_ exhibited in response to the 3 induction protocols.

### 2.8 Development of emulators for the duration and types of atrial arrhythmias

Global sensitivity analysis was also employed to characterise and evaluate how fibrosis and variations in plasma electrolyte concentrations can alter (i) the duration (T_term_) of atrial arrhythmias and (ii) the endpoint and the phenotype. For both T_term_ and the classification of the endpoint, we implemented emulators and fitted the surrogate models using the library PyMC [41].

#### Arrhythmia duration

As noted, in this study we sought to identify arrhythmia terminations within a defined time window: the first 10 seconds after induction. However, each self-sustained arrhythmia can only be characterised by a duration that is longer than 10 seconds. Accordingly, T_term_ is a non-negative, right-censored variable with a three-component mixed distribution. For initiated arrhythmias, T_term_ follows a right-skewed continuous distribution, with right-censoring at t=8900 ms for sustained arrhythmias; non-initiated arrhythmias, conversely, concentrate as ‘mass points’ at T_term_=0. Thus, a unique classical Gaussian process is not suitable as a surrogate for T_term_ (zero floor and right-censoring), because it assumes a unimodal Gaussian observation likelihood. This cannot represent the point mass at zero from non-induced runs, nor can it properly account for right-censored observations at the simulation ceiling. Thus, we utilised a hurdle analysis [27,28]. This was implemented by building the predictive model on two conditionally independent sub-processes: a binary model (hurdle) that predicts if induction (T_term_≥50 ms) occurred; a continuous model (outcome) that models the logarithm of T_term_ only if arrhythmia induction was predicted. The hurdle and the outcome are characterised by the same additive structure: i) a mean-centred 2-D Gaussian process covering the selected range of [Na^+^]_o_ and [K^+^]_o_ values, with Matérn-5/2 covariance kernel and automatic relevance determination (ARD) length-scales; ii) a mean-centred 1-D Gaussian process including the fibrosis amounts, with Matérn-5/2 covariance kernel; and iii) a per-case random effect (hierarchical) with a sum-to-zero constraint ensuring the per-case effects are identifiable with the global intercept by constraining their sum to be exactly zero. The prior probability distributions of the parameters, and the per-case residual, were derived from the cohort data (empirical induction rate, mean and standard deviation of log-duration across and within cases). Finally, a Bernoulli distribution was utilised as the likelihood for the hurdle, while a right-censored normal distribution was utilised as the likelihood for the outcome.

#### Operational definition of the substrate of arrhythmias

The 4 endpoints used to classify the output of these arrhythmia simulations align with their degree of severity (non-induced < terminated < sustained-AT < sustained-AF). However, a standard Gaussian process assuming a continuous likelihood distribution cannot be applied to classification problems having ordinal outcomes. We therefore adopted an ordered-probit model, with a single latent variable that represents the underlying arrhythmia severity. This variable can then be utilised to classify relative probabilities in terms of three ordered ‘cut points’ (the latent variable has the same additive structure adopted in the hurdle model for T_term_). The three cut point values can map the latent variable into one of the 4 ordered classes; the first cut point was placed freely (Normal prior), and the subsequent cut points were parameterised and fitted as positive increments from the previous one, thus enforcing ordering. Finally, an ordered probit distribution was utilised to define the likelihood for the outcome.

When seeking to gain insights into each phenotype, we fitted emulators using the No-U-Turn Sampler (NUTS) algorithm, employing 4 chains with 4096 samples each, discarding the first 2048 (burn-in). We chose a target acceptance rate of 0.95.

For each of the 14 left atrial digital twins, we evaluated the quantities of interest (T_term_ and the four endpoints) for each combination of [Na^+^]_o_ and [K^+^]_o_. We then averaged each output across the 14 predicted values. Thus, the resulting mean value represents the predicted response of the entire 14-patient cohort at a selected plasma electrolyte condition (see Figs 10 and 11).

## 3 Results

### 3.1 PRR in isolated human atrial myocytes

Our initial studies focused on identifying and illustrating the fundamental properties of PRR in (i) a healthy isolated human atrial myocyte and (ii) this atrial myocyte after being modified to exhibit some characteristics of fibrosis (see Methods). Fig 2 consists of plots PRR values for healthy and fibrotic atrial myocytes obtained at four different cycle lengths and following selected variation of electrolyte levels ([Na^+^]_o_, [K^+^]_o_, and [Ca^2+^]_o_) one at a time.

**Fig 2.**
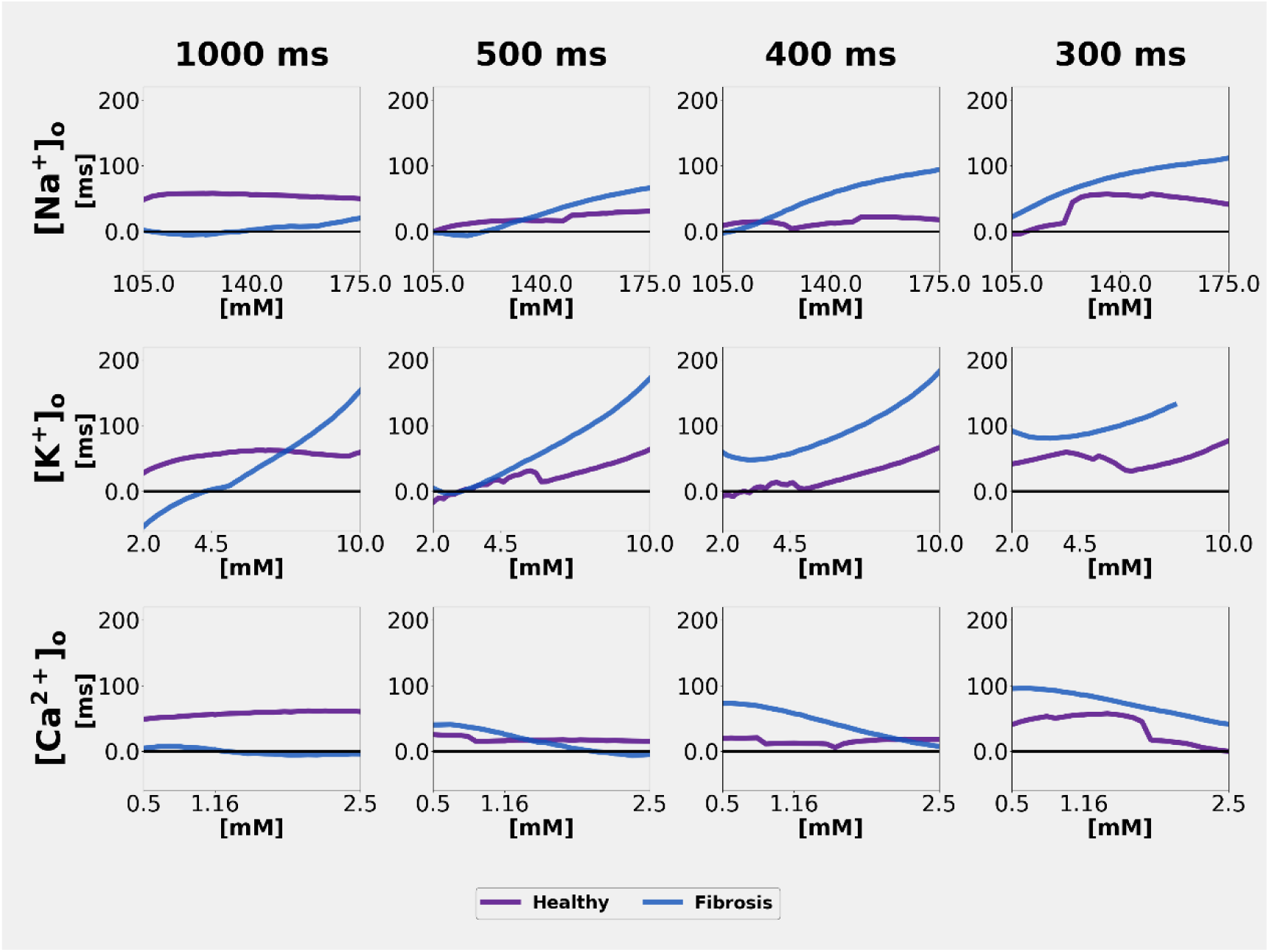
Human atrial single myocyte post-repolarisation refractoriness. Each panel shows post-repolarisation refractoriness (PRR) values in the healthy (purple) and fibrotic (blue) human atrial myocytes. PRR values were obtained at steady state in response to: (i) individual changes in plasma Na^+^, K^+^ or Ca^2+^ values and (ii) a train of stimuli at four different cycle lengths (CL), 1000 ms, 500 ms, 400 ms and 300 ms; (iii) the assumption of healthy or normal conditions (purple) or fibrotic myocyte (blue). When a stimulus train of CL 400 or 300 ms was applied, the membrane APD sometimes exhibited ‘alternans’, and/or intermittent failure to respond to the train of stimuli, characterised as 2:1 capture. When alternans occurred, we computed PRR at the longer APD. The complex dynamic of 2:1, conversely, cannot be analysed using conventional PRR operational definition and therefore it is not shown.

As shown in the middle row of Fig 2, in the healthy atrial myocyte, PRR values (denoted in purple) were weakly sensitive to variations (2-10 mM) in [K^+^]_o_. These changes are presumably produced by the shifts (depolarisation) in the RMP (Figs 3 and 4) and thus the consequent inactivation of the fast inward Na^+^ current INa. Interestingly, at a CL of 1000 ms hypokalaemia ([K^+^]_o_ reductions from e.g. 4.0 to 2.5 mM) reduces PRR values. In contrast, at a CL of either 500 or 400 ms hyperkalaemia tends to lengthen PRR. Conversely, in the setting of fibrosis, PRR values (shown in blue) changed substantially, in response to all variations in [K^+^]_o_.

**Fig 3.**
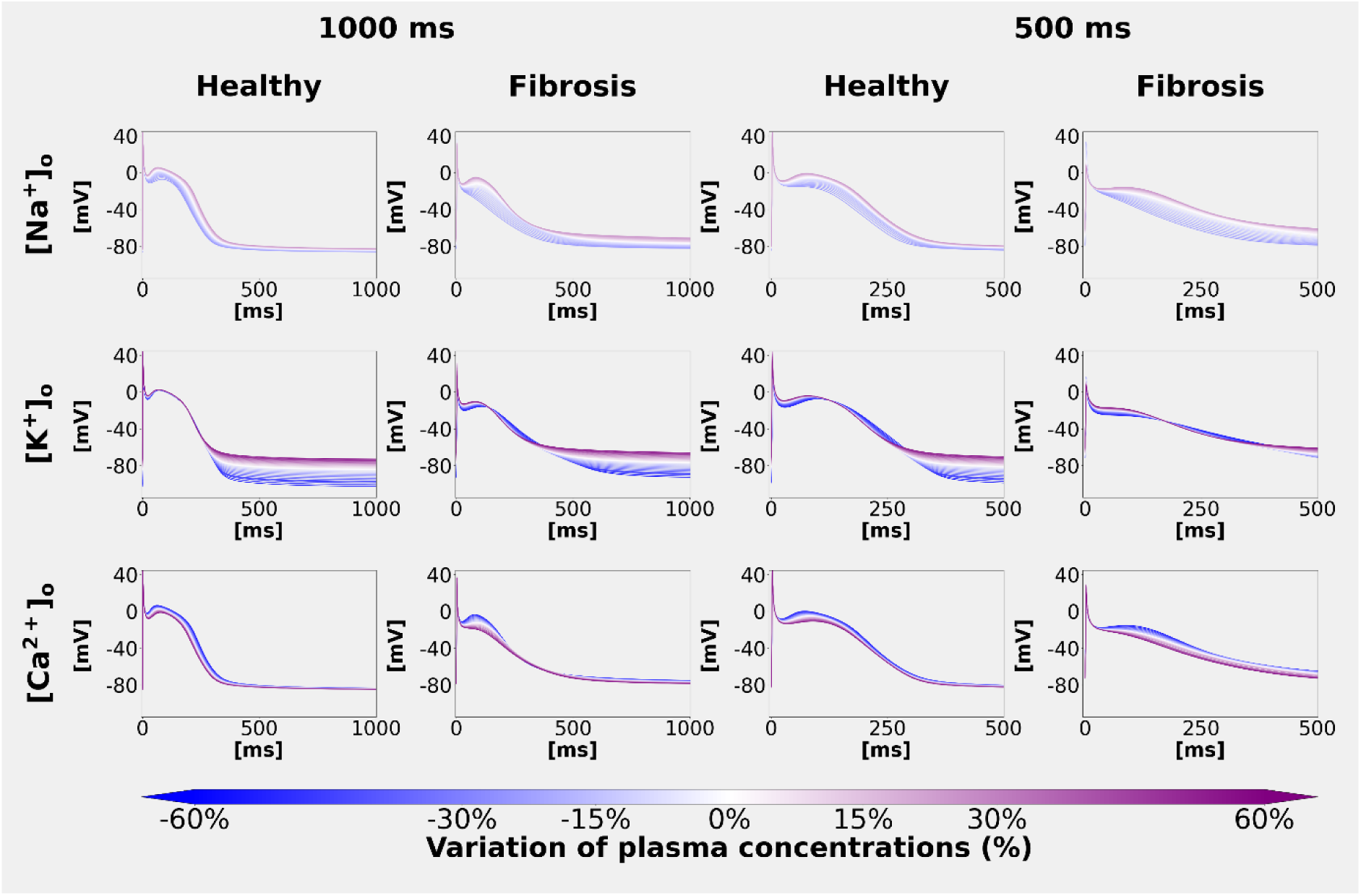
Single human atrial myocyte action potential waveforms recorded following our selected variations in plasma electrolyte concentrations. Human atrial myocyte APs were computed at steady-state at two different CL (CL=1000 ms, left columns, and CL = 500 ms, right columns). The superimposed APs in columns 1 and 3 summarise findings obtained for healthy atrial myocytes, and the data in columns 2 and 4 were obtained from myocytes altered by simulated fibrosis (see Methods). Note the changes in resting membrane potential in rows 1 and 2. The plasma concentration of Na^+^ was varied ±25% from the control baseline (140 mM), and [K^+^]_o_ and [Ca^2+^]_o_ were varied ±60% from the control or baseline values (4.5 mM and 1.16 mM, respectively).

**Fig 4.**
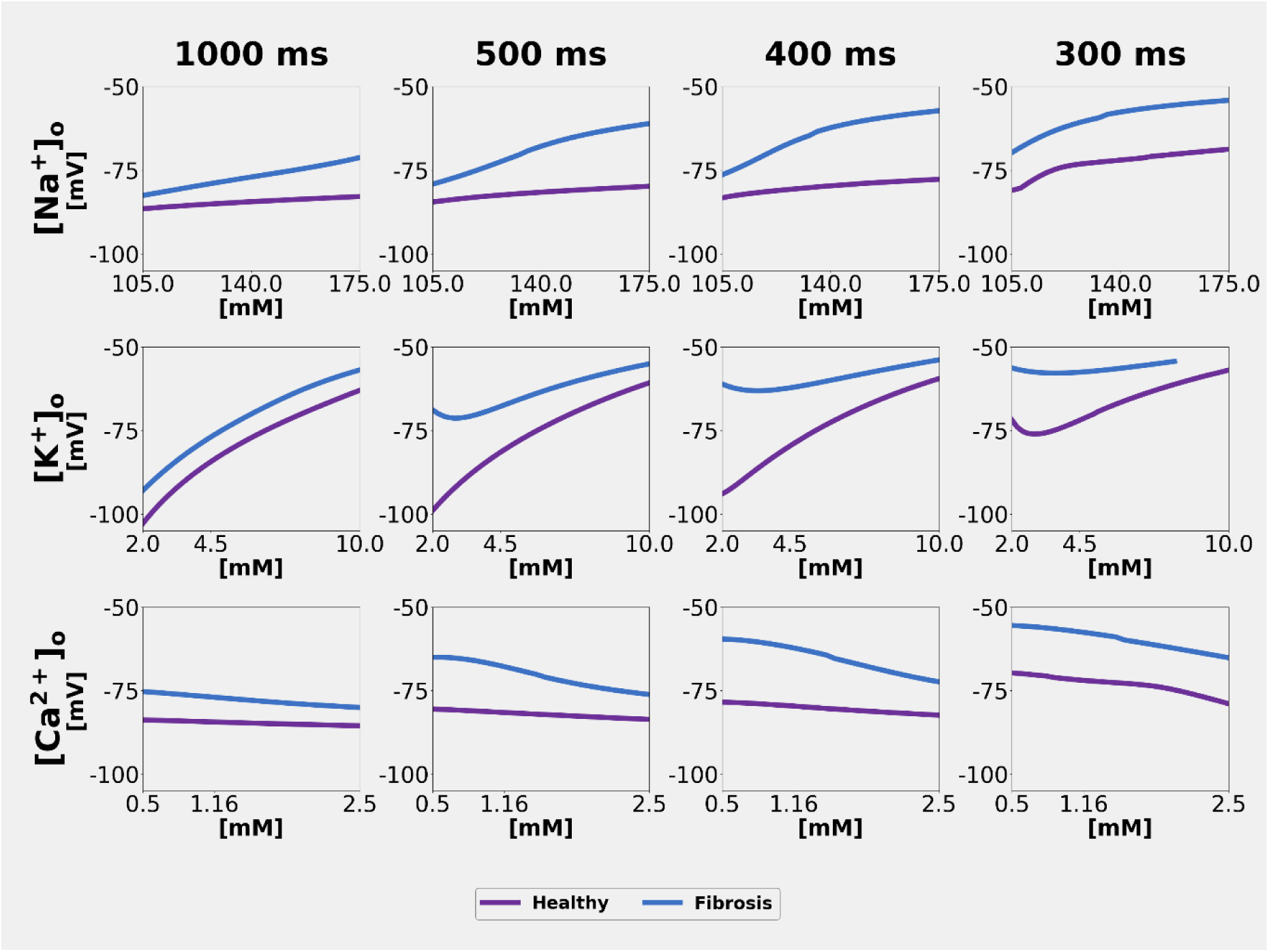
Electrolyte-induced changes in single atrial myocyte resting membrane potential. Illustration of the extent to which individual plasma electrolyte-induced changes can alter the resting membrane potential (RMP) in the healthy and fibrotic human atrial myocytes. RMP values were computed in response to two consecutive stimuli applied at a CL of 1000, 500, 400 and 300 ms (healthy myocyte, purple; fibrotic myocyte, cyan).

Row 1 of Fig 2 also shows that in healthy atrial myocytes variations in [Na^+^]_o_ at a CL of 500 ms and shorter result in small changes in PRR, and these changes are enhanced in the setting of fibrosis. Within the range of alterations in [Ca^2+^]_o_ selected for this study, only very small effects on the shape of the action potential, and hence on PRR, were observed.

As shown in Figs 3 and 4, these descriptive electrolyte-induced changes in human atrial PRR can be understood in terms of (i) the coincident changes in the resting membrane potential (RMP); and (ii) the changes in action potential waveform, including APD. Note that when [K^+^]_o_ is altered the RMP values change significantly, while APD remains quite constant at both CL 1000 ms and CL 500 ms. In contrast, when [Na^+^]_o_ is changed, RMP remains close to baseline values, but the AP waveform tends to be altered (broadened). In the setting of fibrosis combined with hypernatraemia or hypokalaemia, the repolarisation phase of the AP is slowed significantly.

In mammalian atria, it is well-known that even small changes ( <u>±</u>5mV) in the resting potential can significantly alter myocyte and tissue excitability, refractoriness and antiarrhythmic drug sensitivity [12]. Mainly for this reason, our analysis included detailed examinations of RMP in response to the selected plasma electrolyte changes. Our findings are summarised in Fig 4. In summary: in the healthy atrial myocyte only [K^+^]_o_ changes altered RMP, and these effects were substantial (∼35 mV in total). In contrast, in the setting of fibrosis, changes in [Na^+^]_o_ also produced consistent but smaller (± 20 mV changes; see Discussion).

To gain further insight into the relative functional importance of these plasma electrolyte-induced changes in PRR in healthy or fibrotic single atrial myocytes, we next estimated the global sensitivity of PRR to each independent variation of plasma concentrations. These results are summarised in Fig 5A. Again, PRR values for both healthy (the two left-hand plots) and fibrotic (the two right-hand plots) myocytes are shown. This analysis was done at a cycle length of 500 ms, since PRR values at this rate were alternans-free and consistently exhibited a marked sensitivity to plasma electrolyte variations.

**Fig 5.**
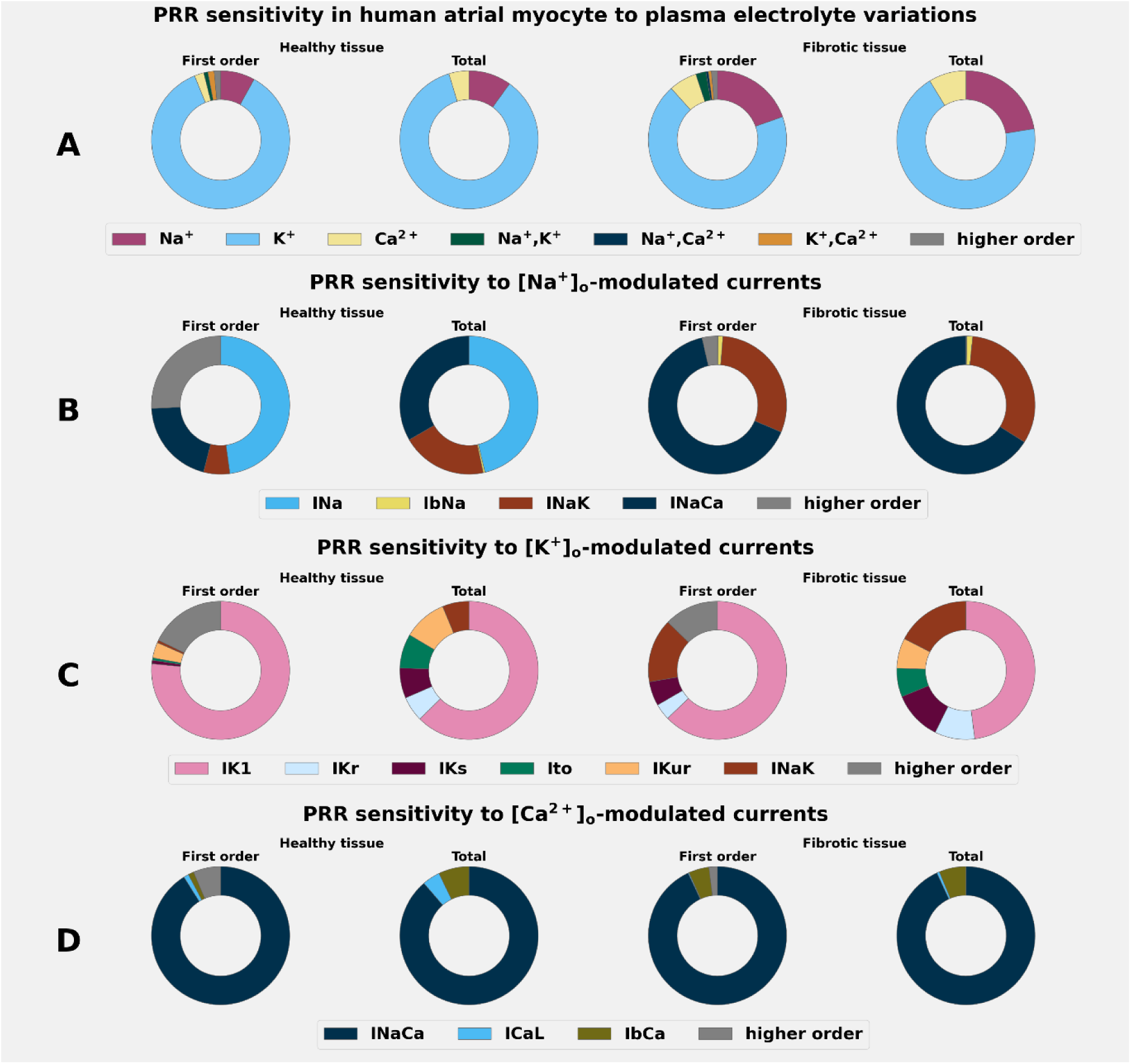
Sensitivity of PRR in human atrial myocyte to selected plasma electrolyte variations and to electrolyte-modulated ionic currents. Row A illustrates the PRR global sensitivity, estimated at a cycle length of 500 ms, and expressed in terms of independent variations in plasma electrolyte concentrations of Na^+^, K^+^, and Ca^2+^, as well as the combinations of Na^+^ and K^+^, Na^+^ and Ca^2+^, as well as K^+^ and Ca^2+^. The PRR global sensitivity to ion-selective channels modulated by Na^+^ (Row B), K^+^ (Row C), and Ca^2+^ (Row D) are also shown. This global sensitivity analysis was applied using healthy atrial myocytes (2 left-hand columns) and in the setting of atrial fibrosis (2 right-hand columns).

The Sobol plots in Rows B, C and D of Fig 5 summarise the GSA analysis of plasma electrolyte changes now expressed in terms of sensitivities to the Na^+^, K^+^ and Ca^2+^-mediated transmembrane ionic currents that modulate the resting potential and the membrane action potential in the human atrial myocyte.

In summary, variations in the selected [K^+^]_o_ alter PRR mainly by changing the inward rectifier potassium current (IK1), both in the healthy myocyte and in the setting of fibrosis. It is known that IK1 affects the resting membrane potential, thus strongly modulating the number of channels available to support the fast Na^+^ inward current (INa) [22]. In the setting of fibrosis, the Sobol plots also reveal a small contribution of the Na^+^/K^+^ pump outward electrogenic current (INaK) to the [K^+^]_o_-driven response. This analysis also revealed that the sensitivity to variations in [Na^+^]_o_ can be accounted for by changes in the Na^+^/Ca^2+^ exchanger (INaCa) and, secondarily, INaK. In contrast, direct contributions of INa were not detectable. The healthy myocyte also showed a minimal sensitivity to variations in [Na^+^]_o_; however, in this case, the predominant channel was the fast [Na^+^] inward current (INa).

### 3.2 Electrolyte-induced changes in PRR and conduction velocity in a human atrial tissue strand

To begin to address some aspects of the translational significance of our findings, the electrolyte-induced changes in PRR in atrial tissue were examined in a series of simulations using the human atrial strand preparation described in the Methods. Results from these studies yielded important functional information concerning electrolyte effects on (i) the propagated as opposed to membrane action potential, (ii) its refractoriness properties, and (iii) conduction velocity changes.

The quite complex patterns of results summarised in Fig 6 show PRR values obtained from a healthy human atrial tissue strand (purple) and in the setting of fibrosis (blue), in response to the same plasma electrolyte changes that were assessed in Fig 2. Data sets were obtained at three basic cycle lengths (CL = 1000, 500, 400 ms), and variations in [Na^+^]_o_ (first row), [K^+^]_o_ (second row), and [Ca^2+^]_o_ (third row) were each studied. Two consistent effects are notable. First, changes in [K^+^]_o_ again result in significant PRR increases. Second, in the setting of fibrosis, the most prominent effect is that at stimulus frequencies ≥ 1Hz there is ‘drop out’ or failure to capture in this atrial tissue strand configuration. In some cases, APD alternans were sometimes observed, but that was not studied in detail.

**Fig 6.**
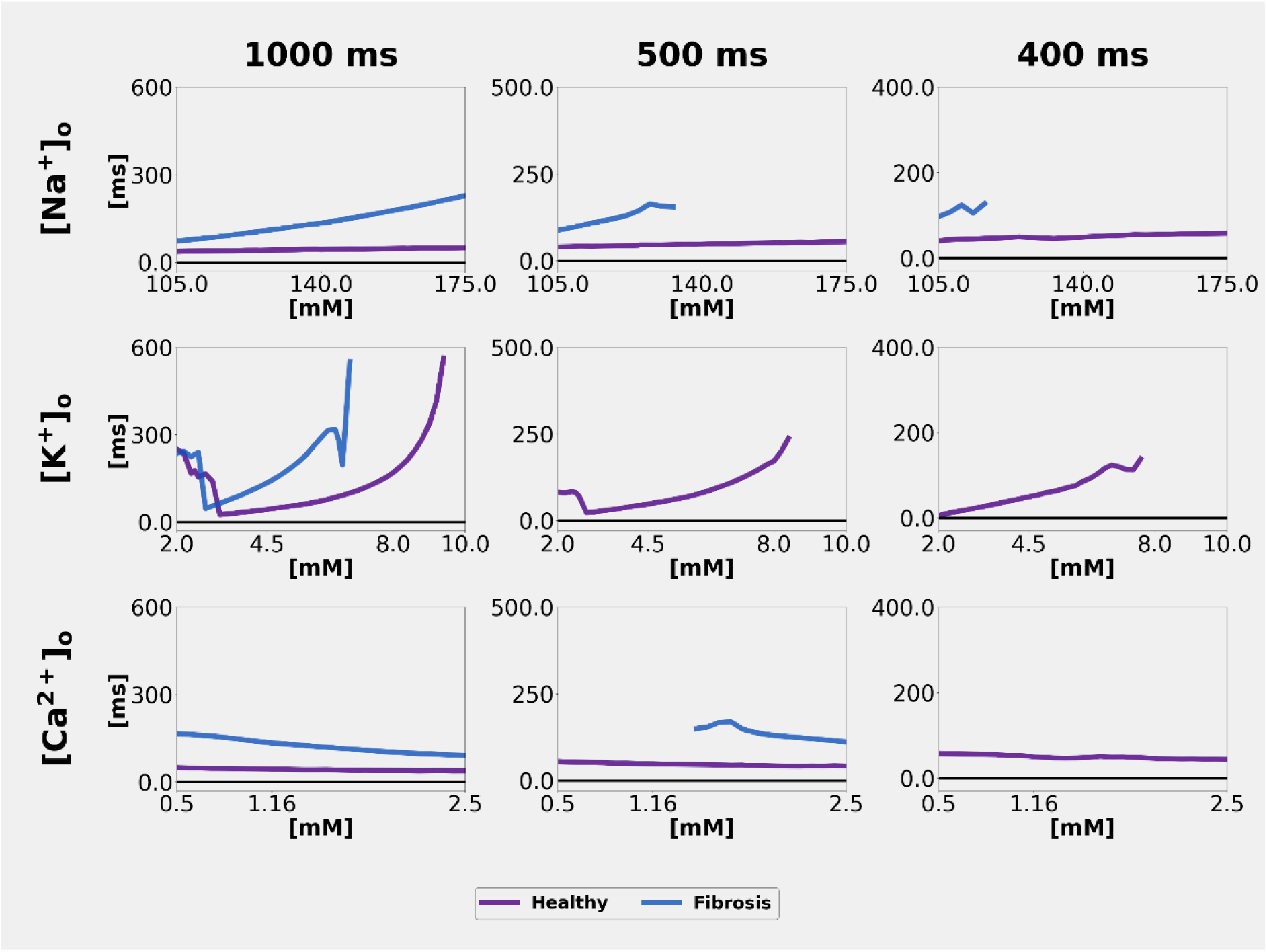
Plasma electrolyte-induced changes in post-repolarisation refractoriness in a human atrial tissue strand. Summary of PRR values in the healthy (purple) and fibrotic (blue) human atrial tissue strands in response to changes in plasma Na^+^, K^+^ or Ca^2+^ values. PRR was computed at CLs of 1000, 500, and 400 ms.

These results in atrial tissue were analysed further by (i) computing the corresponding RMP and AP waveforms (as shown in Fig 7) and (ii) measuring the effects of plasma electrolyte changes on conduction velocity and transmembrane ionic currents (as shown in Figs 8 and 9, respectively). Three types of [K^+^]_o_-induced changes are noteworthy: (i) hypokalaemia significantly hyperpolarises the RMP; (ii) the combination of hyponatraemia and fibrosis hyperpolarises the RMP and markedly reduces the excitability or initial depolarisation phase of the AP; (iii) in the setting of fibrosis, the broadened AP waveform is modulated by changes in [Na^+^]_o_ or [Ca^2+^]_o_, and these two effects are more prominent at a CL of 500 ms than at a CL of 1000 ms.

**Fig 7.**
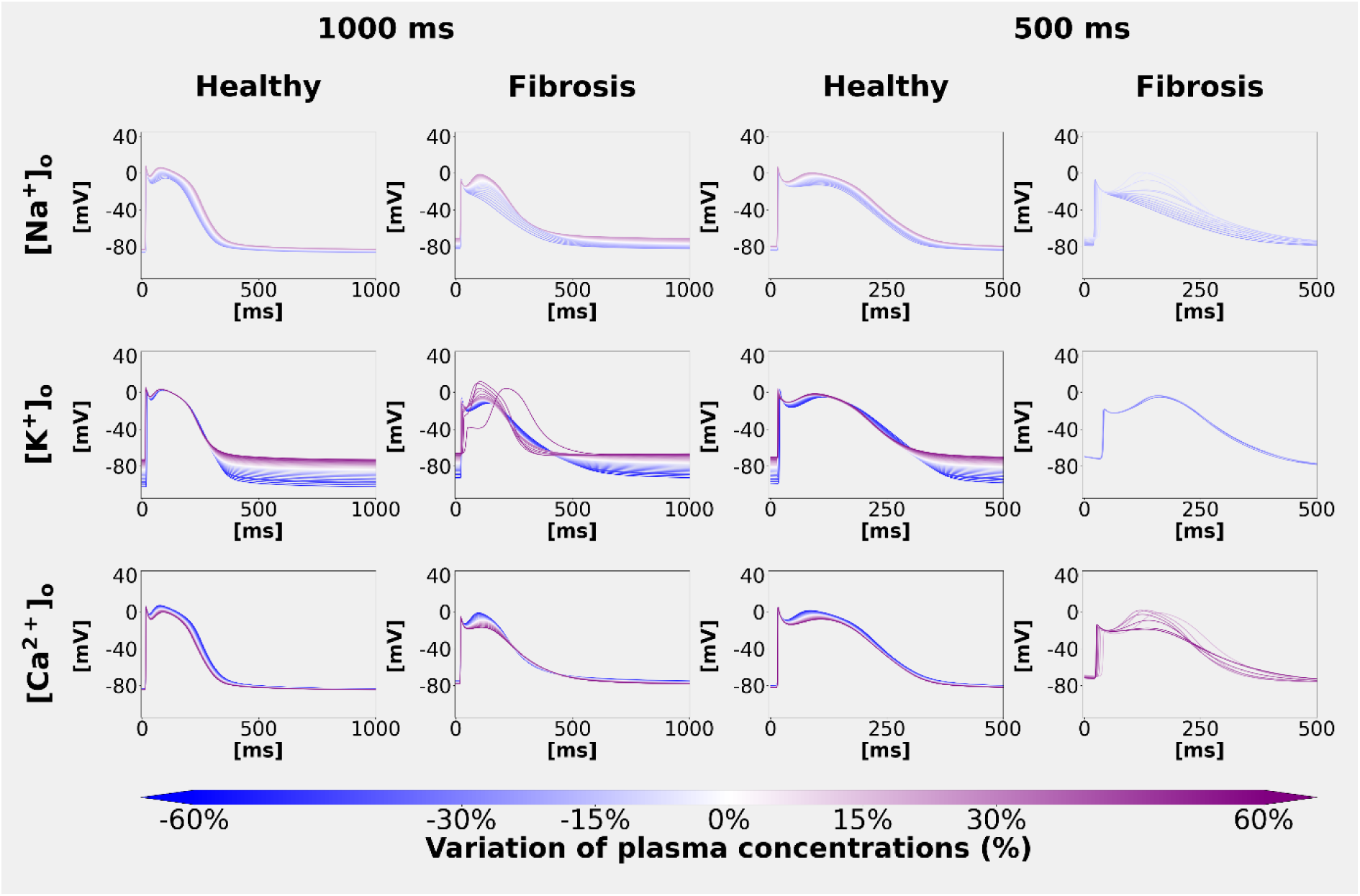
Action potential waveforms recorded at the central section of the atrial tissue strand following our selected variations in plasma electrolyte concentrations. Effects of changes in plasma electrolytes on the resting membrane potential (RMP) and the propagated action potential waveforms measured from the centre (x_2_=5mm) of a human atrial tissue strand (See Methods and text in Results for additional details).

**Fig 8.**
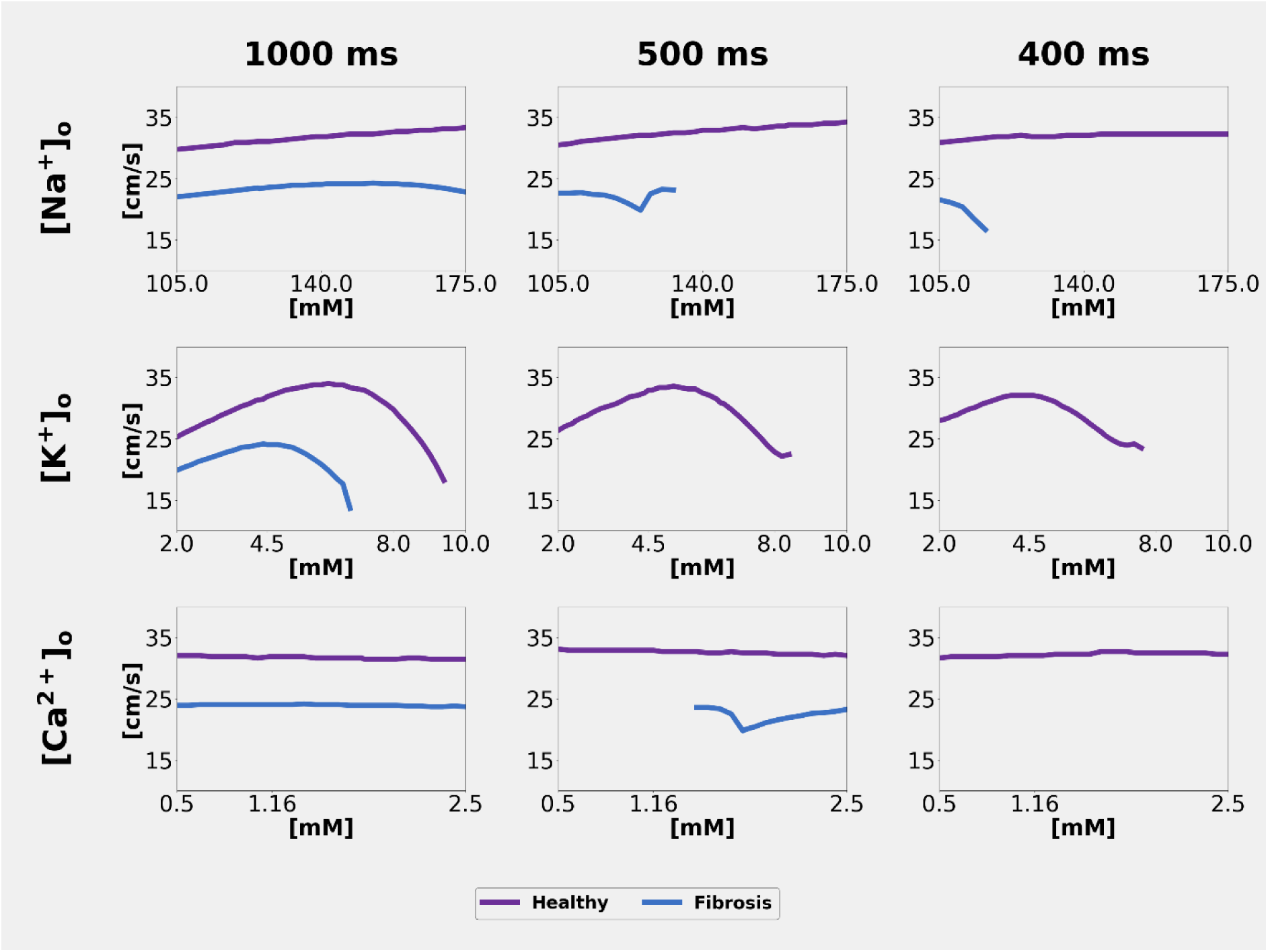
Plasma electrolyte-induced changes in Conduction Velocity in a human atrial tissue strand. Each panel shows CV values in the healthy (purple) and fibrotic (blue) human atrial tissue strands. CV was computed at a CL of 1000, 500, and 400 ms.

**Fig 9.**
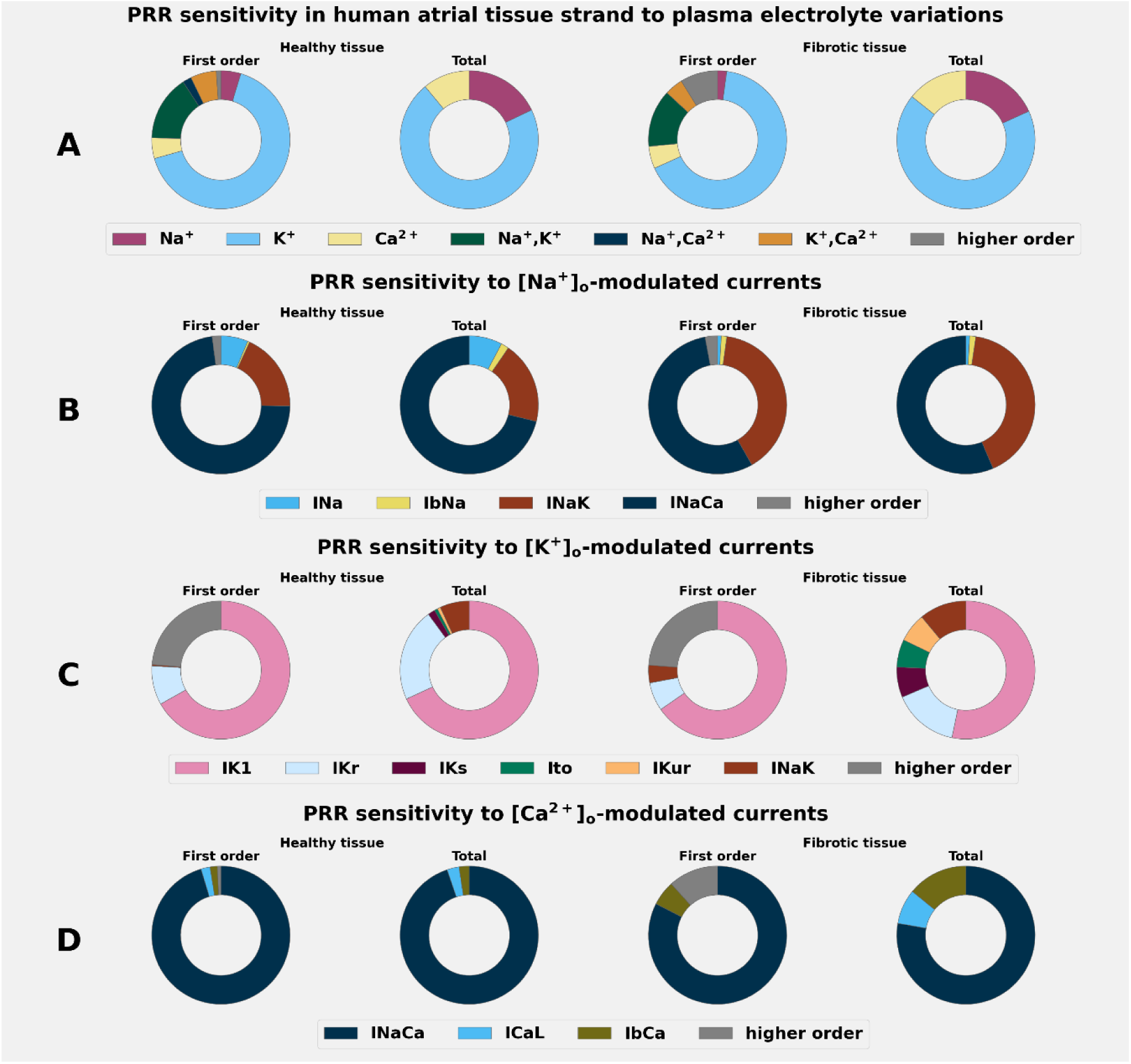
Sensitivity of PRR in human atrial tissue strand to selected plasma electrolyte variations and to electrolyte-modulated ionic currents. Row A illustrates the PRR global sensitivity, estimated at a cycle length of 1000 ms over a human atrial tissue strand 1 cm long, and expressed in terms of independent variations in plasma electrolyte concentrations of Na^+^, K^+^, and Ca^2+^, as well as the combinations of Na^+^ and K^+^, Na^+^ and Ca^2+^, as well as K^+^ and Ca^2+^. The PRR global sensitivity to ion-selective channels modulated by Na^+^ (Row B), K^+^ (Row C), and Ca^2+^ (Row D) is also shown. This global sensitivity analysis was evaluated using healthy atrial tissue strands (2 left columns) and in the setting of atrial fibrosis (2 right columns).

The effects of plasma electrolyte changes on CV of the propagating AP are shown in Fig 8. As expected for mammalian atrial tissue [42], these results clearly show that CV is strongly sensitive to even small variations in [K^+^]_o_, both in healthy tissue (purple) and in the setting of fibrosis (blue). Note also that, as shown in the middle row, in fibrotic tissue, 2:1 conduction block was observed across the entire [K⁺]ₒ plasma concentration range at a stimulus CL ≤ 500 ms. An additional important feature of these tests in healthy atrial strands included, during severe hypokalaemia ([K^+^]_o_ ≤ 3.0 mM), a larger stimulus was required to elicit a regenerative AP (see Discussion).

We next determined the global sensitivity of atrial strand PRR to independent variations of plasma concentrations of electrolytes, as shown in Fig 9A. In addition, GSA was applied to the Na^+^, K^+^ and Ca^2+^ permeant ion channels that underlie RMP resting potential and generate the atrial AP (Fig 9B, C, and D). These procedures were applied to healthy atrial tissue (the two left-hand columns) and in the setting of fibrosis (the two right-hand columns). This analysis of the human atrial strand was performed only at a CL of 1000 ms, since at this rate the substrate is alternans-free. Our findings again clearly showed that even small alterations in [K^+^]_o_ modulate PRR in both healthy and fibrotic conditions (Fig 9A). More detailed inspection of these results at the single transmembrane current level (Fig 9 B, C, D) confirmed that IK1 is the primary mediator in both settings. We also noted that [Na^+^]_o_ - sensitive currents (Fig 9B) revealed a different sensitivity profile between isolated healthy atrial myocytes and healthy atrial tissue strands; isolated atrial myocyte PRR was mainly sensitive to INa, followed by INaCa. The atrial tissue preparation, conversely, revealed a predominant sensitivity to INaCa, followed by INaK, with INa contributing only marginally. In fibrotic tissue, both preparations presented a strong sensitivity to the Na^+^/Ca^2+^ exchanger, followed by the Na^+^/K^+^ pump (INaK). The isolated atrial myocyte and the atrial tissue strand presented similar sensitivities to [Ca^2+^]_o_-modulated currents, dominated by INaCa; ICaL and IbCa contributed only marginally.

This difference reflects the cell-to-cell propagation, in which the Na^+^/Ca^2+^ exchanger modulates the action potential plateau and propagation dynamics more strongly than the direct effect of [Na⁺]ₒ on the INa driving force (see discussions).

### 3.3 Sensitivity of arrhythmia inducibility and sustainability to alterations in plasma electrolytes in human atria

This study was completed by assessing the ways in which changes in plasma electrolytes can induce or sustain atrial arrhythmias. The data in Fig 10B summarise the main finding expressed in terms of arrhythmia duration (T_term_) being modulated by variations in concentrations of [K^+^]_o_ and/or [Na^+^]_o_, based on data obtained from the cohort of 14 left atrial digital twins. The red isolines in this figure denote the probability that an arrhythmia is induced and does not terminate within the simulated window (10 seconds, see Methods). Note that at [K^+^]_o_< 3 mM, hypokalaemia produces longer T_term_ values, i.e. exhibits the highest probability of producing sustained arrhythmias, and that mild hypernatraemia appears to partially compensate the pro-arrhythmic effect of severe hypokalaemia. This analysis also suggests that interactions between hypokalaemia and hyponatraemia result in a longer T_term_.

**Fig 10.**
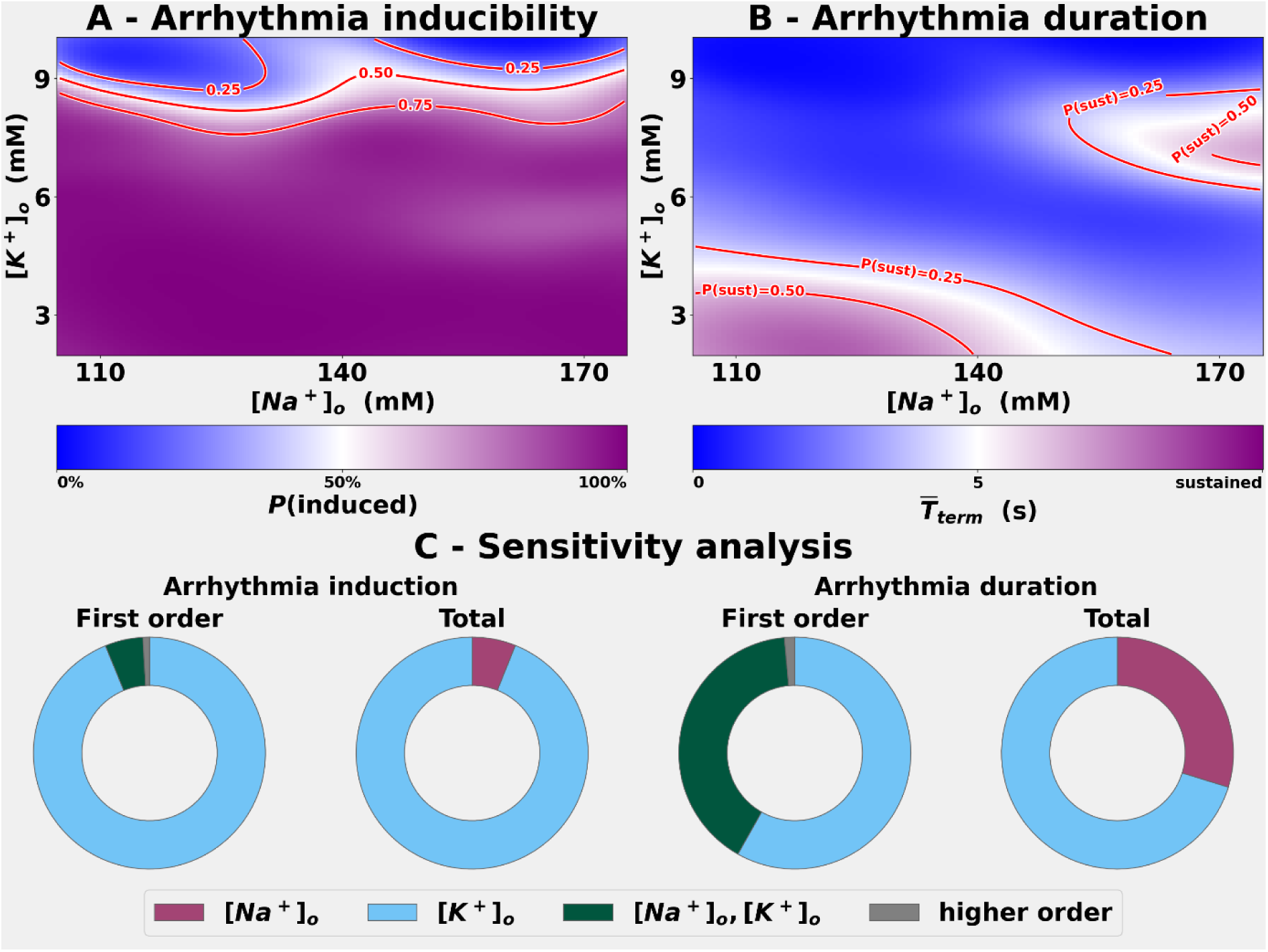
Inducibility and duration of atrial arrhythmias and global sensitivity analysis. Panel A summarises the probability of inducing atrial arrhythmia for the cohort of 14 left atrial digital twins. The red isolines show the (25, 50, 75)% probability of inducing an arrhythmia. Note that changes in [K^+^]_o_ have a significant effect on the probability of inducing an arrhythmia, while the contribution of [Na^+^]_o_ is modest. Panel B shows the duration (T_term_) of the arrhythmia as a function of [Na^+^]_o_ and/or [K^+^]_o_, for the cohort of 14 left atrial digital twins. The red isolines show the probability that an arrhythmia is induced and does not terminate within the simulated window. Panel C shows the sensitivity of inducing an arrhythmia in the human atrium to alterations in [Na^+^]_o_ and/or [K^+^]_o_ (left); and the sensitivity of the duration of the atrial arrhythmia as a function of alterations in [Na^+^]_o_ and/or [K^+^]_o_ (right). Note that inducibility prevalently depends on [K^+^]_o_. Note also that the second-order interactions ([Na^+^]_o_, [K^+^]_o_) reveal that the duration sensitivity to [Na^+^]_o_ depends on the interactions with changes in [K^+^]_o_.

These results also revealed that during hyperkalaemia, induced arrhythmias were of short duration (Panel B) and had a low probability of developing a sustained arrhythmia (Panel A). However, when severe hypernatraemia is also present, T_term_ tended to increase, and the probability of developing a sustained arrhythmia was enhanced. This scenario (simultaneous presence of severe hyperkalaemia and severe hypernatraemia) is rare, although it has been reported [43]. In summary, alterations in [K^+^]_o_ modulate T_term_, while the effect of alterations in [Na^+^]_o_ is dependent on the co-incident value of [K^+^]_o_. This insight is supported by the sensitivity analysis shown in Fig 10C, where [Na^+^]_o_ has second-order effects only.

### 3.4 Probability of atrial arrhythmia induction, termination or endpoint sensitivity to alterations in plasma electrolytes

Fig 11A shows the probability of developing a sustained atrial arrhythmia (AT or AF) as a function of changes in both [Na^+^]_o_ and [K^+^]_o_ for the entire or combined cohort of 14 left atrial digital twins (see Methods). The red isolines correspond to 25, 50 and 75% probability levels. Fig 11B shows the expected endpoint, estimated with the formula: E[Y]= Σ_k_ k·P(Y=k| [Na^+^]_o_, [K^+^]_o_) and as a function of the [Na^+^]_o_ and [K^+^]_o_ levels. The red isolines represent values of E[Y] corresponding to self-terminating arrhythmia (1.0), atrial tachycardia (2.0), or intermediate values between non-induction and self-terminating (0.5) between self-terminating and AT(1.5), and between AT and AF (2.5) where the predicted class distribution is evenly split between adjacent severities; for example, between AT and AF at E[Y]=2.5.

**Fig 11.**
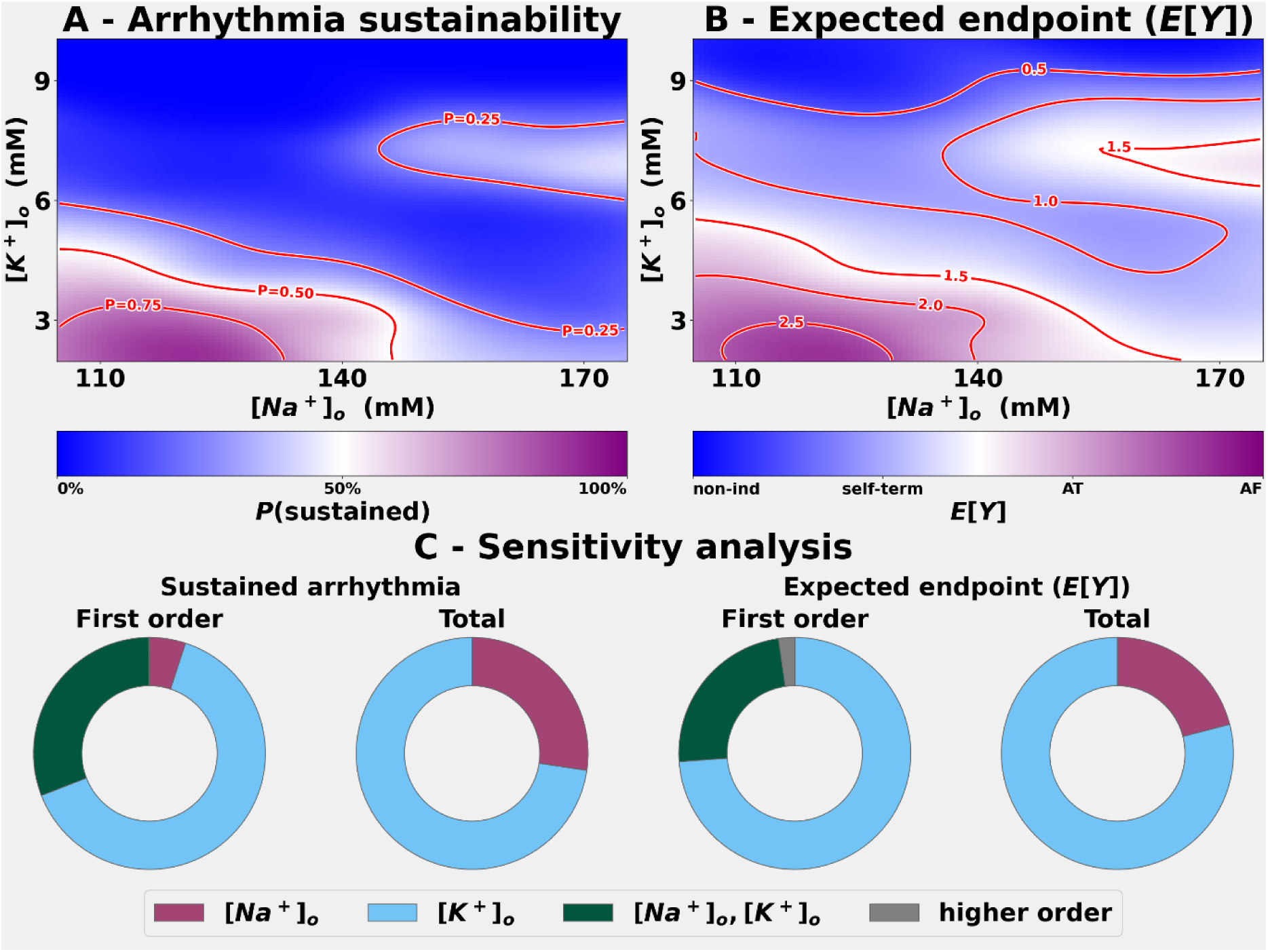
Probability of developing a sustained arrhythmia and expected endpoint and global sensitivity analysis. Panel A shows the probability of developing a sustained atrial arrhythmia (AT or AF) as a function of [Na^+^]_o_ and/or [K^+^]_o_ levels, for the cohort of 14 left atrial digital twins. Red isolines are the 25%, 50%, and 75% probability values. Panel B shows the expected endpoint E[Y] = Σ_k_ k·P(Y=k| [Na^+^]_o_, [K^+^]_o_), as a function of [Na^+^]_o_ and/or [K^+^]_o_ for the same cohort. Red isolines mark half-integer E[Y] levels, indicating where the predicted class distribution is evenly split between adjacent severities (see text for further explanation). Panel C shows the sensitivity of the human atria to develop a sustained arrhythmia to alterations in [Na^+^]_o_ and/or [K^+^]_o_ (left); and the sensitivity of the expected endpoint, evaluated as E[Y] = Σ_k_ k·P(Y=k| [Na^+^]_o_, [K^+^]_o_), to alterations in [Na^+^]_o_ and/or [K^+^] (right). In both indices, the sensitivity to [Na^+^]_o_ was strongly dictated by second-order interactions ([Na^+^]_o_, [K^+^]_o_).

In summary, when the data in Figs 10 and 11 are considered together, these findings reveal a pattern: the atrial arrhythmia endpoint is strongly modulated by [K^+^]_o_. The small influence of changes in [Na^+^]_o_ is noteworthy only when these changes occur jointly with [K^+^]_o_ changes (second-order effects only, as shown in Fig 11C). Significant hypokalaemia makes the atrial substrate strongly pro-arrhythmic and prone to generate AF in both baseline and fibrotic settings. The combined effects of significant hyperkalaemia and severe hypernatraemia can also produce a pro-arrhythmic substrate. However, AT is the most prominent sustained arrhythmia.

## 4 Discussion

### Main findings

The main finding from this digital twin-based study of the effects of changes in plasma electrolyte on human atrial post-repolarisation refractoriness (PRR) is that even a quite small change in [K^+^]_o_ can be the dominant determinant of PRR in both the isolated myocyte and the human atrial tissue strand. These changes in [K^+^]_o_ act primarily by regulating the resting membrane potential (RMP). This is caused by alterations in the nonlinear background K^+^ current, IK1 [11]. In mammalian atria, even very small changes in RMP regulate the fast-inward Na^+^ current, INa, through its steady-state inactivation [12]. As shown in Figs 2 and 6, even modest depolarisation of RMP of mild hyperkalaemia can markedly lengthen ERP, and therefore increase PRR [13]. Other studies have shown that lengthening PRR can thus enhance the antiarrhythmic action [16] of sodium-channel blockers [17,18].

Our results also provide insights into some of how electrolyte changes, combined with fibrosis, can alter PRR in the human atrial substrate. In brief, the fibrosis-dependent decrease in the magnitude of the Na^+^ and K^+^ currents lower excitability, decreases both conduction velocity and safety factor for conduction and tend to depolarise the myocyte/atrial substrate resting potential (see Limitations).

### Distinctions between electrolyte effects on membrane vs propagated atrial action potentials and their refractoriness

We note also that the multi-scale approach that we have chosen for this study has revealed significant differences when PRR analysis is applied to the single atrial myocyte vs the atrial strand preparation. These differences can be understood in terms of classical biophysical models for (i) the membrane action potential (the single myocyte) versus (ii) the regenerative propagated action potential that is present in the syncytium – the human atrial strand. These principles and their differences have been clearly identified and described in classical analysis of action potential initiation and conduction in excitable tissue [44]. That work, and more recent analyses of the Safety Factor for conduction [45] identify the complexities of both the transmembrane current flow and capacitive discharge as they interact to yield to the net currents required for either excitation or repolarisation when AP propagation occurs.

These findings appear to be augmented or enhanced in the setting of atrial fibrosis. The way in which we have modelled fibrosis (reductions in the ion channel mediated conductances for Na^+^, Ca^2+^, and K^+^) results in the substrate having an increased input resistance, reduced conduction velocity, and much less net transmembrane current for either depolarization or repolarization. Not surprisingly, therefore, both ERP and PRR are changed with the consistent indication that the combination of fibrosis and electrolyte changes may result in PRR being meaningful in clinical settings where fibrosis is present or suspected.

As shown in Figs 2 and 6, PRR was only weakly sensitive to [Na^+^]_o_, and this effect was produced by different transmembrane currents in each setting. In the healthy myocyte, this effect was mainly due to INa. In the single atrial myocyte, in presence of the low-excitability state produced by the fibrotic remodelling described above, INa could no longer produce a full-amplitude upstroke; consequently, the contribution of INa to the sensitivity of PRR to [Na^+^]_o_ became small. Here, the GSA analysis favoured excitability by the Na⁺-dependent transporters, the Na^+^/Ca^2+^ exchanger (INaCa) and the Na^+^/K^+^ pump (INaK). In contrast, in the healthy human atrial tissue strands the sensitivity to [Na^+^]_o_ acted through INaCa, whereas under fibrosis INaCa and INaK had equal weight. These differences follow from the distinction drawn above: the myocyte receives an applied stimulus that always fires a depolarisation unless the cell is in absolute refractoriness, whereas in the strand the cells are depolarised by the propagation front. Under the latter conditions [Na^+^]_o_ influences ERP primarily through INaCa, by altering its electrochemical driving force during the plateau and thus modifying the plateau voltage, the APD, and consequently the diastolic interval available for INa recovery in downstream tissue, [46]. Direct recordings in human atrial trabeculae support this coupling between INaCa, [Na^+^]_o_ and the plateau state [47]. Although the effects of INaCa on the action potential waveform are present in isolated-cell simulations, their consequences for propagation through downstream tissue depend on electrotonic coupling between neighbouring cells. Where propagation safety rather than excitability is limiting, repolarisation-reserve currents such as Ito and IKur become more important through their effect on recovery in the adjacent myocytes [45].

### Insights in relation to anti-arrhythmic drugs or drug combination therapies

These differences between single myocyte or membrane AP vs multi-cellular or atrial strand propagated APs are relevant to anti-arrhythmic drugs action. It is well known that the efficacy of Class-I and Class-III agents in the atrium depends on the resting membrane potential [48,49], on combining sodium-channel block with a modest rise in [K^+^]_o_ [15] and, at IK1, details on the change in the inward rectifying K⁺ current with direct modulation of Kir2.1 [50,51]. Clinically, even the mild electrolyte excursions of routine practice may change atrial PRR far enough to either start or stop AF. Perhaps the best example can be found in reports of hyperkalaemia converting permanent AF to sinus rhythm [14].

As shown in Figs 10 and 11, when we further investigated how the selected electrolyte alterations can affect the type and the duration of the arrhythmias, we concluded that the onset, duration and the phenotype of atrial arrhythmias exhibited a sensitivity to variations in [K^+^]_o_. In general, our findings align with previous reports of there being a significant role for IK1 in re-entrant atrial arrhythmia dynamics [52]. Our findings also suggested that changes in [Na^+^]_o_ modulated conduction velocity (Fig A1 in S1 Appendix), the duration of atrial arrhythmia and the risk or phenotype only via second-order interactions with [K^+^]_o_. For the management of AT or AF, this suggests that alterations in [Na^+^]_o_ carry an added risk of conduction slowing, as well as an effect on refractoriness. There is epidemiological data linking the urinary Na^+^/K^+^ ratio and 24-hour Na^+^ excretion to a higher prevalence of AF [9]. As a final part of this analysis, we examined whether raising [K^+^]_o_ into the high-normal range (4.5-5 mM) has an effect on the human atrium somewhat similar to the anti-arrhythmic effects that have been observed in the human ventricle [53]. As shown in Fig A3 in S1 Appendix, our four-class GP surrogate approach yielded results that supported the POTCAST trial [53] hypothesis that higher extracellular potassium reduces sustained atrial arrhythmias and identified most of the suppressed events as sustained AF. Global sensitivity and 2-D surface analyses (Fig 11) revealed that elevated [K^+^]_o_ was a dominant monotone driver, suggesting that serum-K^+^ management is a plausible, actionable translational consideration for this clinical cohort.

### Limitations

We are aware, however, that both the conclusions that we have made and our suggestions concerning translational applications of our findings have limitations. These include:

1. This analysis, like that in our previous study ([22]) is dependent on the mathematical model for the human atrial resting potential and action potential that we have utilised. The Courtemanche model of the human atrial AP has been evaluated and used previously; and our modification of the ways in which changes in [K^+^]_o_ alter IK1 ensure that simulations of [K^+^]_o_-induced changes on RMP are reliable. However, the Courtemanche model does not include high-fidelity intracellular Ca^2+^ buffering or RyR2-mediated Ca^2+^ release that underlies triggered activity and delayed afterdepolarisations [54,55]. Moreover, although changes in intracellular Na^+^ are accounted for, the Na^+^ ‘accumulation’ during rapid pacing is modest. Accordingly, the rate-dependent modulation of INaK and INaCa may be underestimated.
2. We note that the definition of PRR differs somewhat between published studies. As noted in Methods, in this study, the PRR time period was defined as ERP-APD_75_. Previous literature has set the repolarisation time point between APD_60_-APD_90_ [56,57]. Although, the specific PRR cut point will affect the exact PRR value, it is unlikely to affect our conclusions, which we reproduced in sensitivity analysis across a range of cycle lengths.
3. The differences in the sensitivity to changes in plasma electrolytes (mainly [K^+^]_o_ and [Na^+^]_o_) between membrane and conducted action potentials is an important insight. However, some features of the descriptors for the conducted action potential model (as shown in Fig 7) e.g. the intercellular resistance and the maximum conductance for INa) are model dependent and could influence both the Safety Factor for regenerative conduction and the ‘liminal length’ of our strand [58].
4. As noted in Methods, the specific way in which fibrosis is modelled in this and previous studies, although computationally tractable, has limitations. Additional analyses should include mathematical descriptions of the fibroblasts (i) being a separate cell type in the atrial syncytium, and (ii) expressing a fibroblast specific complement of ion channels e.g. Na^+^, K^+^ and TRP.
5. Our consideration of the effects of changes in plasma Ca^2+^, [Ca^2+^]_o_, does not include the well-known surface charge screening effects on the voltage-dependence of the Na^+^, K^+^ and Ca^2+^ conductances [59]. As we have acknowledged previously [22], this improvement should be implemented, particularly given the well-known stabilising or anti-arrhythmic effects of increasing [Mg^2+^]_o_ in mammalian atria, including humans [60].

## 5 Conclusions

This study identified variations in plasma [K^+^]_o_ as the principal determinant in altering human atrial post-repolarisation refractoriness (PRR) in computational studies combining an isolated human atrial myocyte, tissue strand, and patient-specific left atrial digital twins. Severe hypokalaemia rendered the substrate strongly pro-arrhythmic, while maintaining plasma [K^+^]_o_ in the high-normal range (4.5 to 5 mM) reduced the burden of atrial arrhythmias. Hence, plasma [K^+^]_o_ management emerges as a plausible, actionable handle on atrial arrhythmia risk, one that now warrants prospective clinical confirmation.

## Acknowledgements

All simulations used the ARCHER2 UK National Supercomputing Service.

## Data availability

All data required to reproduce the results presented here were previously published and are publicly available in the Zenodo repository (https://doi.org/10.5281/zenodo.5801337) [24]. No restrictions exist on data availability.

The analysis code (myocyte and tissue sensitivity analysis, arrhythmia analysis) is deposited in a public GitHub repository: https://github.com/cesare-corrado/PRR-human-atrium-arrhythmias.

## 6 Supporting information

**S1 Appendix. Supplementary methods and figures.** Global sensitivity analysis of conduction velocity, sub-cohort selection criteria, termination time and organisation-index definitions, and mirroring of the POTCAST study (Sections A1-A5, Figs A1-A3).

## Notes

### Competing Interest Statement

The authors have declared no competing interest.

## References

[1] S.S. Chugh, R. Havmoeller, K. Narayanan, D. Singh, M. Rienstra, E.J. Benjamin, R.F. Gillum, Y.H. Kim, J.H. McAnulty, Z.J. Zheng, M.H. Forouzanfar, M. Naghavi, G.A. Mensah, M. Ezzati, C.J.L. Murray, Worldwide epidemiology of atrial fibrillation: A global burden of disease 2010 study, Circulation 129 (2014) 837–847. 10.1161/CIRCULATIONAHA.113.005119.

[2] S. Cheng, J.Z. He, Y. Han, S. Han, P. Li, H. Liao, J. Guo, Global burden of atrial fibrillation/atrial flutter and its attributable risk factors from 1990 to 2021, Europace 26 (2024) euae195. 10.1093/europace/euae195.

[3] K.T.S. Konings, C.J.H.J. Kirchhof, J.R.L.M. Smeets, H.J.J. Wellens, O.C. Penn, M.A. Allessie, High-density mapping of electrically induced atrial fibrillation in humans, Circulation 89 (1994) 1665–1680. 10.1161/01.CIR.89.4.1665.

[4] P. Kirchhof, A.J. Camm, A. Goette, A. Brandes, L. Eckardt, A. Elvan, T. Fetsch, C.I. van Gelder, D. Haase, M. Haegeli Laurent, F. Hamann, H. Heidbüchel, G. Hindricks, J. Kautzner, K.-H. Kuck, L. Mont, G.A. Ng, J. Rekosz, N. Schoen, U. Schotten, A. Suling, J. Taggeselle, S. Themistoclakis, E. Vettorazzi, P. Vardas, K. Wegscheider, S. Willems, M.J.G. Crijns Harry, G. Breithardt, Early rhythm-control therapy in patients with atrial fibrillation, N. Engl. J. Med. 383 (2020) 1305–1316. 10.1056/NEJMoa2019422.

[5] P.L. Rensma, M.A. Allessie, W.J.E.P. Lammers, F.I.M. Bonke, M.J. Schalij, Length of excitation wave and susceptibility to reentrant atrial arrhythmias in normal conscious dogs, Circ. Res. 62 (1988) 395–410. 10.1161/01.RES.62.2.395.

[6] M.C.E.F. Wijffels, C.J.H.J. Kirchhof, R. Dorland, M.A. Allessie, Atrial fibrillation begets atrial fibrillation: A study in awake chronically instrumented goats, Circulation 92 (1995) 1954–1968. 10.1161/01.CIR.92.7.1954.

[7] B.J. Paice, K.R. Paterson, F. Onyanga-Omara, T. Donnelly, J.M.B. Gray, D.H. Lawson, Record linkage study of hypokalaemia in hospitalized patients, Postgrad. Med. J. 62 (1986) 187–191. 10.1136/pgmj.62.725.187.

[8] L.M. Einhorn, M. Zhan, V.D. Hsu, L.D. Walker, M.F. Moen, S.L. Seliger, M.R. Weir, J.C. Fink, The frequency of hyperkalemia and its significance in chronic kidney disease, Arch. Intern. Med. 169 (2009) 1156–1162. 10.1001/archinternmed.2009.132.

[9] S. Tokioka, N. Nakaya, R. Hatanaka, K. Nakaya, M. Kogure, I. Chiba, M. Takase, K. Nochioka, K. Susukita, H. Metoki, T. Nakamura, M. Ishikuro, T. Obara, Y. Hamanaka, M. Orui, T. Kobayashi, A. Uruno, E.N. Kodama, S. Nagaie, S. Ogishima, Y. Izumi, N. Fuse, S. Kuriyama, S. Yasuda, A. Hozawa, Association between sodium- and potassium-related urinary markers and the prevalence of atrial fibrillation, Circ. J. 89 (2025) 757–764. 10.1253/circj.CJ-24-0780.

[10] A. Burashnikov, M. Pourrier, J.K. Gibson, J.J. Lynch, C. Antzelevitch, Rate-dependent effects of vernakalant in the isolated non-remodeled canine left atria are primarily due to block of the sodium channel comparison with ranolazine and dl-sotalol, Circ. Arrhythm. Electrophysiol. 5 (2012) 400–408. 10.1161/CIRCEP.111.968305.

[11] M.A.G. Van Der Heyden, T. Jespersen, Pharmacological exploration of the resting membrane potential reserve: Impact on atrial fibrillation, Eur. J. Pharmacol. 771 (2016) 56–64. 10.1016/j.ejphar.2015.11.026.

[12] L. Skibsbye, T. Jespersen, T. Christ, M.M. Maleckar, J. van den Brink, P. Tavi, J.T. Koivumäki, Refractoriness in human atria: Time and voltage dependence of sodium channel availability, J. Mol. Cell. Cardiol. 101 (2016) 26–34. 10.1016/J.YJMCC.2016.10.009.

[13] S. V. Pandit, M. Warren, S. Mironov, E.G. Tolkacheva, J. Kalifa, O. Berenfeld, J. Jalife, Mechanisms underlying the antifibrillatory action of hyperkalemia in guinea pig hearts, Biophys. J. 98 (2010) 2091–2101. 10.1016/j.bpj.2010.02.011.

[14] E. Romeo, M. D’Alto, G. Santarpia, G. Di Nardo, B. Sarubbi, M.G. Russo, R. Calabrò, Hyperkalemia-induced conversion of permanent atrial fibrillation to normal sinus rhythm: A case report, J. Cardiovasc. Med. 12 (2011) 678–680. 10.2459/JCM.0b013e32833ce871.

[15] A. Burashnikov, C. Antzelevitch, Mild elevation of extracellular potassium greatly potentiates the effect of sodium channel block to cardiovert atrial fibrillation: The Lankenau approach, Heart Rhythm 20 (2023) 1257–1264. 10.1016/j.hrthm.2023.05.005.

[16] M.R. Franz, R.A. Gray, P. Karasik, H.J. Moore, S.N. Singh, Drug-induced post-repolarization refractoriness as an antiarrhythmic principle and its underlying mechanism, Europace 16 (2014) iv39–iv45. 10.1093/europace/euu274.

[17] K. Fukuda, J. Watanabe, T. Yagi, Y. Wakayama, M. Nakano, M. Kondo, K. Kumagai, M. Miura, K. Shirato, H. Shimokawa, A sodium channel blocker, pilsicainide, produces atrial post-repolarization refractoriness through the reduction of sodium channel availability, Tohoku J. Exp. Med. 225 (2011) 35–42. 10.1620/tjem.225.35.

[18] P. Milberg, G. Frommeyer, S. Ghezelbash, S. Rajamani, N. Osada, R. Razvan, L. Belardinelli, G. Breithardt, L. Eckardt, Sodium channel block by ranolazine in an experimental model of stretch-related atrial fibrillation: Prolongation of interatrial conduction time and increase in post-repolarization refractoriness, Europace 15 (2013) 761–769. 10.1093/europace/eus399.

[19] M. Courtemanche, R.J. Ramirez, S. Nattel, Ionic mechanisms underlying human atrial action potential properties: insights from a mathematical model, Am. J. Physiol. Heart Circ. Physiol. 275 (1998) H301–H321. 10.1152/ajpheart.1998.275.1.H301.

[20] S. Severi, C. Corsi, E. Cerbai, From in vivo plasma composition to in vitro cardiac electrophysiology and in silico virtual heart: The extracellular calcium enigma, Philos. Trans. R. Soc. A Math. Phys. Eng. Sci. 367 (2009) 2203–2223. 10.1098/rsta.2009.0032.

[21] M. Clerx, G.R. Mirams, A.J. Rogers, S.M. Narayan, W.R. Giles, Immediate and delayed response of simulated human atrial myocytes to clinically-relevant hypokalemia, Front. Physiol. 12 (2021) 651162. 10.3389/fphys.2021.651162.

[22] C. Corrado, C.H. Roney, S.M. Narayan, W.R. Giles, S.A. Niederer, The effect of clinically relevant changes in extracellular electrolyte concentrations on human atrial arrhythmias, Commun. Med. 6 (2026) 7. 10.1038/s43856-025-01260-4.

[23] C.H. Roney, A. Pashaei, M. Meo, R. Dubois, P.M. Boyle, N.A. Trayanova, H. Cochet, S.A. Niederer, E.J. Vigmond, Universal atrial coordinates applied to visualisation, registration and construction of patient specific meshes, Med. Image Anal. 55 (2019) 65–75. 10.1016/j.media.2019.04.004.

[24] C. Roney, I. Sim, J. Yu, M. Beach, A. Mehta, J. Solis-Lemus, I. Kotadia, J. Whitaker, C. Corrado, O. Razeghi, E. Vigmond, S. Narayan, M. O’Neill, S. Williams, S. Niederer, Predicting atrial fibrillation recurrence by combining population data and virtual cohorts of patient-specific left atrial models [Data set], (2022). 10.5281/zenodo.5801337.

[25] A. Saltelli, P. Annoni, I. Azzini, F. Campolongo, M. Ratto, S. Tarantola, Variance based sensitivity analysis of model output. Design and estimator for the total sensitivity index, Comput. Phys. Commun. 181 (2010) 259–270. 10.1016/j.cpc.2009.09.018.

[26] S. Longobardi, A. Sher, S.A. Niederer, In silico identification of potential calcium dynamics and sarcomere targets for recovering left ventricular function in rat heart failure with preserved ejection fraction, PLoS Comput. Biol. 17 (2021) e1009646. 10.1371/JOURNAL.PCBI.1009646.

[27] J. Mullahy, Specification and testing of some modified count data models, J. Econom. 33 (1986) 341–365. 10.1016/0304-4076(86)90002-3.

[28] J.G. Cragg, Some statistical models for limited dependent variables with application to the demand for durable goods, Econometrica 39 (1971) 829–844. 10.2307/1909582.

[29] G. Plank, A. Loewe, A. Neic, C. Augustin, Y.L. Huang, M.A.F. Gsell, E. Karabelas, M. Nothstein, A.J. Prassl, J. Sánchez, G. Seemann, E.J. Vigmond, The openCARP simulation environment for cardiac electrophysiology, Comput. Methods Programs Biomed. 208 (2021) 106223. 10.1016/j.cmpb.2021.106223.

[30] H. Gelband, H.L. Bush, M.R. Rosen, R.J. Myerburg, B.F. Hoffman, Electrophysiologic properties of isolated preparations of human atrial myocardium., Circ. Res. 30 (1972) 293–300. 10.1161/01.RES.30.3.293.

[31] J. Keener, J. Sneyd, Mathematical Physiology, Springer Cham, 2026. 10.1007/978-3-031-83217-8.

[32] J. Herman, W. Usher, SALib: An open-source Python library for Sensitivity Analysis, J. Open Source Softw. 2 (2017) 97. 10.21105/joss.00097.

[33] W. Schroeder, K. Martin, B. Lorensen, The Visualization Toolkit (4th ed.), Kitware, 2006.

[34] C.H. Roney, I. Sim, J. Yu, M. Beach, A. Mehta, J. Alonso Solis-Lemus, I. Kotadia, J. Whitaker, C. Corrado, O. Razeghi, Predicting atrial fibrillation recurrence by combining population data and virtual cohorts of patient-specific left atrial models, Circ. Arrhythm. Electrophysiol. 15 (2022) e010253. 10.1161/CIRCEP.121.010253.

[35] C.H. Roney, J.D. Bayer, S. Zahid, M. Meo, P.M.J. Boyle, N.A. Trayanova, M. Haïssaguerre, R. Dubois, H. Cochet, E.J. Vigmond, Modelling methodology of atrial fibrosis affects rotor dynamics and electrograms, Europace 18 (2016) iv146–iv155. 10.1093/europace/euw365.

[36] C.H. Roney, M.L. Beach, A.M. Mehta, I. Sim, C. Corrado, R. Bendikas, J.A. Solis-Lemus, O. Razeghi, J. Whitaker, L. O’Neill, G. Plank, E. Vigmond, S.E. Williams, M.D. O’Neill, S.A. Niederer, In silico comparison of left atrial ablation techniques that target the anatomical, structural, and electrical substrates of atrial fibrillation, Front. Physiol. 11 (2020). 10.3389/fphys.2020.572874.

[37] E. Matene, V. Jacquemet, Fully automated initiation of simulated episodes of atrial arrhythmias, Europace 14 (2012) v17–v24. 10.1093/europace/eus271.

[38] P.M. Boyle, T. Zghaib, S. Zahid, R.L. Ali, D. Deng, W.H. Franceschi, J.B. Hakim, M.J. Murphy, A. Prakosa, S.L. Zimmerman, Computationally guided personalized targeted ablation of persistent atrial fibrillation, Nat. Biomed. Eng. 3 (2019) 870–879. 10.1038/s41551-019-0437-9.

[39] J. Tuan, F. Osman, M. Jeilan, S. Kundu, R. Mantravadi, P.J. Stafford, G.A. Ng, Increase in organization index predicts atrial fibrillation termination with flecainide post-ablation: Spectral analysis of intracardiac electrograms, Europace 12 (2010) 488–493. 10.1093/europace/eup405.

[40] Y. Takahashi, P. Sanders, P. Jaïs, M. Hocini, R. Dubois, M. Rotter, T. Rostock, C.J. Nalliah, F. Sacher, J. Clémenty, M. Haïssaguerre, Organization of frequency spectra of atrial fibrillation: Relevance to radiofrequency catheter ablation, J. Cardiovasc. Electrophysiol. 17 (2006) 382–388. 10.1111/j.1540-8167.2005.00414.x.

[41] O. Abril-Pla, V. Andreani, C. Carroll, L. Dong, C.J. Fonnesbeck, M. Kochurov, R. Kumar, J. Lao, C.C. Luhmann, O.A. Martin, M. Osthege, R. Vieira, T. Wiecki, R. Zinkov, PyMC: a modern, and comprehensive probabilistic programming framework in Python, PeerJ Comput. Sci. 9 (2023) e1516. 10.7717/peerj-cs.1516.

[42] A. Nygren, W.R. Giles, Mathematical simulation of slowing of cardiac conduction velocity by elevated extracellular [K+] in a human atrial strand, Ann. Biomed. Eng. 28 (2000) 951–957. 10.1114/1.1308489.

[43] J. Jacobson, D. Bohn, Severe hypernatremic dehydration and hyperkalemia in an infant with gastroenteritis secondary to rotavirus, Ann. Emerg. Med. 22 (1993) 1360–1362. 10.1016/S0196-0644(05)81270-3.

[44] J.J.B. Jack, D. Noble, R.W. Tsien, Electric Current Flow in Excitable Cells, Clarendon Press, 1975.

[45] P.M. Boyle, W.H. Franceschi, M. Constantin, C. Hawks, T. Desplantez, N.A. Trayanova, E.J. Vigmond, New insights on the cardiac safety factor: Unraveling the relationship between conduction velocity and robustness of propagation, J. Mol. Cell. Cardiol. 128 (2019) 117–128. 10.1016/j.yjmcc.2019.01.010.

[46] D.L. Campbell, W.R. Giles, K. Robinson, E.F. Shibata, Studies of the sodium-calcium exchanger in bull-frog atrial myocytes., J. Physiol. 403 (1988) 317–340. 10.1113/jphysiol.1988.sp017251.

[47] U. Schotten, S. de Haan, S. Verheule, E.G.A. Harks, D. Frechen, E. Bodewig, M. Greiser, R. Ram, J. Maessen, M. Kelm, M. Allessie, D.R. Van Wagoner, Blockade of atrial-specific K+-currents increases atrial but not ventricular contractility by enhancing reverse mode Na+/Ca2+-exchange, Cardiovasc. Res. 73 (2007) 37. 10.1016/j.cardiores.2006.11.024.

[48] F. Syeda, A.P. Holmes, T.Y. Yu, S. Tull, S.M. Kuhlmann, D. Pavlovic, D. Betney, G. Riley, J.P. Kucera, F. Jousset, J.R. de Groot, S. Rohr, N.A. Brown, L. Fabritz, P. Kirchhof, PITX2 modulates atrial membrane potential and the antiarrhythmic effects of sodium-channel blockers, J. Am. Coll. Cardiol. 68 (2016) 1881–1894. 10.1016/j.jacc.2016.07.766.

[49] A.P. Holmes, P. Saxena, S.N. Kabir, C. O’Shea, S.M. Kuhlmann, S. Gupta, D. Fobian, C. Apicella, M. O’Reilly, F. Syeda, J.S. Reyat, G.L. Smith, A.J. Workman, D. Pavlovic, L. Fabritz, P. Kirchhof, Atrial resting membrane potential confers sodium current sensitivity to propafenone, flecainide and dronedarone, Heart Rhythm 18 (2021) 1212–1220. 10.1016/J.HRTHM.2021.03.016.

[50] S.F. Noujaim, J.A. Stuckey, D. Ponce-Balbuena, T. Ferrer-Villada, A. López-Izquierdo, S. Pandit, C.J. Calvo, K.R. Grzeda, O. Berenfeld, J.A. Sánchez Chapula, J. Jalife, Specific residues of the cytoplasmic domains of cardiac inward rectifier potassium channels are effective antifibrillatory targets, FASEB J. 24 (2010) 4302–4312. 10.1096/fj.10-163246.

[51] R. Caballero, P. Dolz-Gaitón, R. Gómez, I. Amorós, A. Barana, M.G. De La Fuente, L. Osuna, J. Duarte, A. López-Izquierdo, I. Moraleda, E. Gálvez, J.A. Sánchez-Chapula, J. Tamargo, E. Delpón, Flecainide increases Kir2.1 currents by interacting with cysteine 311, decreasing the polyamine-induced rectification, Proc. Natl. Acad. Sci. U. S. A. 107 (2010) 15631–15636. 10.1073/pnas.1004021107.

[52] S.F. Noujaim, S. V. Pandit, O. Berenfeld, K. Vikstrom, M. Cerrone, S. Mironov, M. Zugermayr, A.N. Lopatin, J. Jalife, Up-regulation of the inward rectifier K+ current (IK1) in the mouse heart accelerates and stabilizes rotors, J. Physiol. 578 (2007) 315–326. 10.1113/jphysiol.2006.121475.

[53] C. Jøns, C. Zheng, U.C.G. Winsløw, E.M. Danielsen, T. Sakthivel, E.A. Frandsen, H. Saffi, S.S. Vakilzadeh-Hashemi, K.J. Haugan, N.E. Bruun, K.K. Iversen, H.S. Bosselmann, N. Risum, H. Bundgaard, Increasing the potassium level in patients at high risk for ventricular arrhythmias, N. Engl. J. Med. 393 (2025) 1979–1989. 10.1056/nejmoa2509542.

[54] J. Heijman, H. Sutanto, H.J.G.M. Crijns, S. Nattel, N.A. Trayanova, Computational models of atrial fibrillation: Achievements, challenges, and perspectives for improving clinical care, Cardiovasc. Res. 117 (2021) 1682–1699. 10.1093/cvr/cvab138.

[55] D. Dobrev, X.H.T. Wehrens, Calcium-mediated cellular triggered activity in atrial fibrillation, J. Physiol. 595 (2017) 4001–4008. 10.1113/JP273048.

[56] P.F. Kirchhof, C.L. Fabritz, M.R. Franz, Postrepolarization refractoriness versus conduction slowing caused by class I antiarrhythmic drugs antiarrhythmic and proarrhythmic effects, Circulation 97 (1998) 2567–2574. 10.1161/01.CIR.97.25.2567.

[57] S.M. Narayan, M.R. Franz, P. Clopton, E.J. Pruvot, D.E. Krummen, Repolarization alternans reveals vulnerability to human atrial fibrillation, Circulation 123 (2011) 2922–2930. 10.1161/CIRCULATIONAHA.110.977827.

[58] D. Noble, The relation of Rushton’s ‘liminal length’ for excitation to the resting and active conductances of excitable cells, J. Physiol. 226 (1972) 573–591. 10.1113/jphysiol.1972.sp009998.

[59] D.L. Campbell, W.R. Giles, J.R. Hume, D. Noble, E.F. Shibata, Reversal potential of the calcium current in bull-frog atrial myocytes., J. Physiol. 403 (1988) 267–286. 10.1113/jphysiol.1988.sp017249.

[60] F. Cacioppo, D. Reisenbauer, H. Herkner, J. Oppenauer, N. Schuetz, J. Niederdoeckl, S. Schnaubelt, S. Gupta, M. Lutnik, A. Simon, A.O. Spiel, N. Buchtele, H. Domanovits, A.N. Laggner, M. Schwameis, Association of intravenous potassium and magnesium administration with spontaneous conversion of atrial fibrillation and atrial flutter in the emergency department, JAMA Netw. Open 5 (2022) e2237234. 10.1001/jamanetworkopen.2022.37234.

